# *map3k1* suppresses terminal differentiation of migratory eye progenitors in planarian regeneration

**DOI:** 10.1101/2024.10.11.617745

**Authors:** Katherine C. Lo, Christian P. Petersen

## Abstract

Proper stem cell targeting and differentiation is necessary for regeneration to succeed. In organisms capable of whole body regeneration, considerable progress has been made identifying wound signals initiating this process, but the mechanisms that control the differentiation of progenitors into mature organs are not fully understood. Using the planarian as a model system, we identify a novel function for *map3k1,* a MAP3K family member possessing both kinase and ubiquitin ligase domains, to negatively regulate terminal differentiation of stem cells during eye regeneration. Inhibition of *map3k1* caused the formation of multiple ectopic eyes within the head, but without controlling overall head, brain, or body patterning. By contrast, other known regulators of planarian eye patterning like *WntA* and *notum* also regulate head regionalization, suggesting *map3k1* acts distinctly. Eye resection and regeneration experiments suggest that unlike Wnt signaling perturbation, *map3k1* inhibition did not shift the target destination of eye formation in the animal. Instead, *map3k1(RNAi)* ectopic eyes emerge in the regions normally occupied by migratory eye progenitors, and the onset of ectopic eyes after *map3k1* inhibition coincides with a reduction to eye progenitor numbers. Furthermore, RNAi dosing experiments indicate that progenitors closer to their normal target are relatively more sensitive to the effects of *map3k1,* implicating this factors in controlling the site of terminal differentiation. Eye phenotypes were also observed after inhibition of *map2k4, map2k7, jnk,* and *p38*, identifying a putative pathway through which *map3k1* prevents differentiation. Together, these results suggest that *map3k1* regulates a novel control point in the eye regeneration pathway which suppresses the terminal differentiation of progenitors during their migration to target destinations.

**Author Summary:** During adult regeneration, progenitors must migrate and differentiate at the proper locations in order to successfully restore lost or damaged organs and tissues, yet the mechanisms underlying these abilities are not fully understood. The planarian eye is a model to study this problem, because this organ is regenerated using migratory progenitors that travel long distances through the body in an undifferentiated state prior to terminal differentiation upon their arrival at target destinations. We determined that a pathway involving the MAP kinase kinase kinase *map3k1* holds planarian eye progenitors in an undifferentiated state during their transit. Inhibition of *map3k1* caused a dramatic body transformation in which migratory progenitors differentiate inappropriately early, and in the wrong locations, into mature eyes. By analyzing this phenotype and measuring the change to eye progenitor abundance after *map3k1* inhibition, we found that *map3k1* prevents ectopic differentiation of eye cells rather than mediating body-wide patterning through the Wnt pathway. Our study argues that whole-body regeneration mechanisms involve separate steps to control patterning and progenitor differentiation.

## Introduction

The process of regeneration restores damaged tissue in a spatially coordinated manner that produces new parts in the correct locations and proportions. This process requires precise control of programs which detect injury, re-establish global body axis, activate stem cell proliferation and differentiation, target progenitor to the proper areas, and ultimately assemble cells into functional tissues and organs. While the pathways controlling progenitor specification for regeneration have become increasingly resolved, still little is known about the processes enabling the targeting of these cells to appropriate locations. The planarian *Schmidtea mediterranea* can replace nearly any adult tissue after injury and are a model for understanding whole-body regeneration mechanisms (1). This ability requires the neoblast population of pluripotent stem cells, which constitute the organism’s only adult somatic proliferative cell type (2, 3). Elimination of neoblasts by gamma-irradiation prevents regeneration, while transplantation of single neoblasts into irradiated animals rescues this regeneration defect (2). After injury, neoblasts proliferate and specify into specialized progenitors in order to differentiate into the different cell types required for building new tissue. The progenitors building regionalized tissues such as eyes and pharynx are specified in broader domains along the body axes and are believed to subsequently migrate to specific target positions in order to build new organs through regeneration or to homeostatically maintain existing organs through gradual cell replacement (4). However, the mechanisms ensuring that stem cells terminally differentiate only at the correct locations are not well understood.

The planarian eye is a paradigm for understanding the mechanisms of organ regeneration. Planarians have two bilateral eyes residing in the anterior part of body, containing populations of *opsin^+^* rhabdomeric photoreceptor neurons (PRNs), as well as *tyrosinase^+^* optic cup cells (also known as pigment cups cells, PCCs). PRNs produce ARRESTIN^+^ axons which project ipsilaterally and contralaterally to enervate to the ventral brain. Axon guidance involves interactions with a specific set of *notum^+^* and *fzd5/8-4^+^* muscle cells which act as guidepost cells to provide cues for the proper formation of the neuronal circuit and visual axon bundles (5). The lateral portion of each eye is composed of PCCs believed to use melanin to allow light shadowing of PRN inputs enabling appropriate left/right discrimination for negative phototaxis (6–8). New eyes can regenerate after surgical eye ablation, damage to the eye regions, or after decapitation during head regeneration. Eye regeneration first involves specification of naïve neoblasts into eye progenitors through expression of transcription factors *ovo, eyes absent (eya),* and *sine oculis-1/2 (six-1/2)*. In uninjured animals, *ovo^+^* eye progenitors are dispersed throughout the anterior half of the body, and in eye regeneration, these cells also abundant within “trails” of cells located posterior and lateral to the newly forming eyes. BrdU pulse-chase experiments suggest these trails of eye progenitors migrate and are the source of new eye cells during regeneration (7). During specification, eye progenitors are partitioned into distinct eye cell subpopulations: *otxA^+^/ovo^+^* progenitors which differentiate into *opsin^+^* PRNs and *sp6-9^+^/dlx^+^/ovo^+^*progenitors that generate *tyrosinase^+^* PCCs (6, 7). The expression of these fate-specifying transcription factors is also retained after eye progenitors terminally differentiate into mature eye cells. Proper formation of each subclass of PRN cells also involves additional eye transcription factors *soxB, meis, klf,* and *foxQ2*, because their inhibition by RNAi resulted in aberrant optic cup phenotypes such as small eyes and elongated optic cup morphology, without decreasing eye progenitor numbers (6). Eye differentiation was reduced following inhibition of *egfr-4* (9) and *bcat-4* (10), and formation of PRNs increased after inhibition of the NuRD complex component *p66* (11). Thus, a complex network of factors regulates specification and terminal differentiation of the eye cells in regeneration.

In addition to controlled eye cell specification, body-wide patterning factors ensure that eyes form in the correct position and proportion. Position control genes (PCGs) are signaling factors expressed in the body-wall muscle which establish global body axis and determine regional tissue identities (12, 13). The anterior-posterior (AP) axis in planarians is broadly defined by a gradient of β-catenin dependent Wnt signaling, with *wnt1* expression defining the posterior and Wnt inhibitors such as *notum* expressed in the anterior (14–17). Inhibition of *notum* in uninjured animals or in regenerating head fragments causes an anterior shift of regional head identity resulting in formation of a set of eyes anterior to the original pre-existing eyes (18). Inhibition of other head patterning factors such as *wntA* (also known as *wnt11-6*), a Wnt signaling factor which regulates brain size via a negative feedback loop with *notum* (19), and *noudarake* (*ndk*), a fibroblast growth factor receptor-like (FGFRL) gene which restricts brain tissues to the head (20, 21), can also cause a set of ectopic eyes to form posterior to pre-existing eyes. Inhibition of *src-1*, a non-receptor tyrosine kinase, caused phenotypes resembling a combination of the Wnt/FGFRL pathways to result in posterior expansion of head, formation of posterior ectopic eyes, as well as ectopic trunk formation (22). Inhibition of *slit* and *wnt5*, regulators of the medial-lateral (ML) axis, cause medial or more lateral ectopic eyes to form, respectively (23). Modification of Wnt/FGFRL signaling and Wnt5/Slit signaling also causes AP and ML changes to brain structure, respectively, suggesting these patterning signals more broadly control head regional identity that also affects the eyes (19–24). Interestingly, although regeneration of new eyes occurs at precise locations defined by PCG genes, eye homeostasis can occur in broader domains. Eyes transplanted into the anterior but not posterior regions of the body are homeostatically maintained by migratory eye progenitors (25), and inhibition of patterning factors can shift the site of new eye regeneration without altering eye homeostasis (18, 22). However, it is not yet clear what signals may coordinate eye progenitors to their target destinations and enable their proper differentiation specifically at the location of existing eyes.

Here we identify a pathway involving *mitogen-activated protein kinase kinase kinase 1* (*map3k1*) that specifically regulates the site of terminal differentiation of migratory eye progenitors. *map3k1* (also known as MEKK1) is a mitogen-associated protein kinase kinase kinase known to transduce extracellular signals into a diverse set of intracellular responses, including cell survival, cell migration, and in apoptosis. MAP3K1 factors contain a C-terminal kinase domain characteristic of MAP kinases, as well as a RING domain with E3 ubiquitin ligase activity (26, 27), a unique combination among MAP3Ks. Our research identifies a novel role for planarian *map3k1* in suppressing terminal differentiation and preserving the eye progenitor state until they reach their target destination.

## Results

### *map3k1* RNAi causes formation of ectopic eyes

We conducted a small-scale RNAi screen of signaling molecules whose expression had been described previously as activated by injury. We identified a single planarian homolog of *map3k1* (dd_Smed_v6_5198_0_1) which encodes a protein with predicted kinase and RING finger domains typical of MAP3K1 family members (Fig S1A). Prior studies in *Schmidtea mediterranea* had identified the *map3k1* transcript as rapidly induced within the first 10 minutes after injury (28, 29) but the role of this gene in regeneration was unclear. In order to investigate the function of this gene, we inhibited *map3k1* by RNAi using 6 dsRNA doses administered over 2 weeks, then challenged the worms to regenerate. *map3k1(RNAi)* animals succeeded at formation of blastemas, but newly formed heads produced ectopic eyes in positions that appeared in lateral-posterior locations within the blastema (Fig 1A). The ectopic eye phenotype had the greatest penetrance in head fragments (15/15 animals, 100% penetrance), compared to regenerating trunks (13/15 animals, 87%) and regenerating tail fragments (9/15, 60%). We used FISH to confirm that ectopic eyes in *map3k1(RNAi)* animals contain both *opsin^+^* PRNs and *tyrosinase^+^* PCCs (Fig 1B), and anti-ARRESTIN antibody staining shows that *map3k1(RNAi)* ectopic eyes can also make axon projections to the brain (Fig 1C). These results suggest that *map3k1(RNAi)* ectopic eyes possess the morphology of normal eyes.

**Fig 1.**
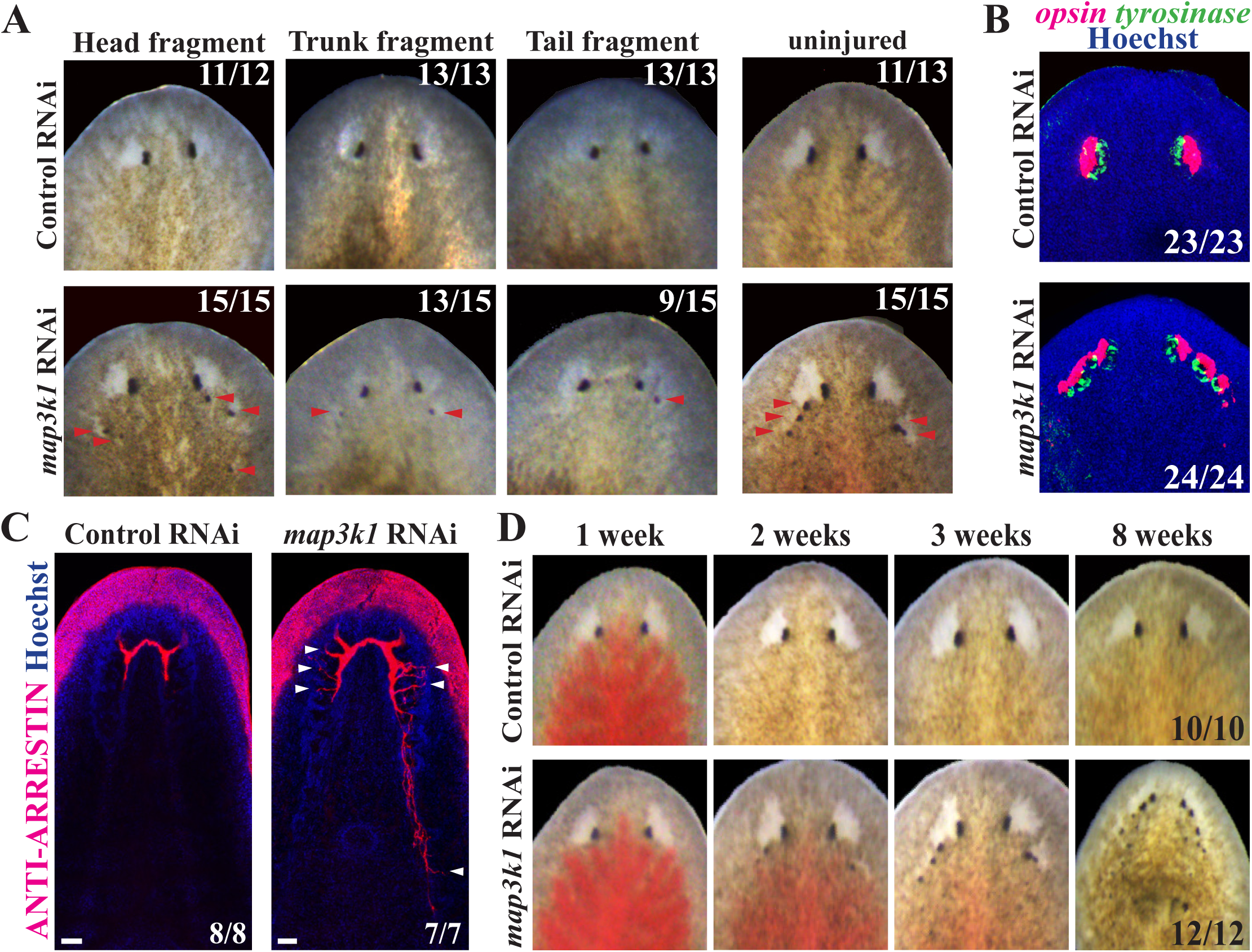
*map3k1* RNAi causes formation of ectopic eyes in regenerating and uninjured planarians. (A-B) Animals were fed control or *map3k1* dsRNA 6 times over 2 weeks, then amputated into head, trunk and tail fragments and allowed to regenerate for 14 days or left uninjured for an equal amount of time, followed by (A) live imaging or (B) FISH to detect *opsin* and *tyrosinase* markers of eye cells. In each type of regenerating fragment, *map3k1* RNAi caused formation of ectopic eyes (red arrowheads) that contained *opsin^+^* photoreceptor neurons and *tyrosinase^+^*pigment cup cells. (C) Homeostatic *map3k1(RNAi)* and control animals were stained with anti-ARRESTIN antibody to mark photoreceptor neuron axons. *map3k1* inhibition caused formation of ectopic ARRESTIN^+^ axons projecting toward the brain (white arrowheads). Scale bars 50μm. (D) Uninjured animals were fed with dsRNA twice per week for the times indicated and live imaged in a timeseries to visualize the progression of the *map3k1(RNAi)* phenotype. *map3k1(RNAi)* caused a progressive formation of additional eyes mainly located within the head region. Number of animals scored for each condition are indicated in the panels.

Given *map3k1*’s demonstrated transcriptional activation in the early wound response, we wanted to determine whether its role in eye patterning required its injury-induced gene expression. To test this possibility, we inhibited *map3k1* homeostatically and examined the impact on eye formation (Fig 1A). Long-term inhibition of *map3k1* in the absence of injury resulted in highly penetrant formation of ectopic eyes (15/15, 100%), indicating *map3k1* regulates an eye formation process common to both regeneration and homeostasis. The *map3k1(RNAi)* phenotype appeared to progress over time, with ectopic eyes appearing by 2 weeks of RNAi, and additional eyes appearing over time through 3-8 weeks of treatment (Figs 1D and S1B). *map3k1(RNAi)* animals survived through at least 8 weeks of RNAi, suggesting that *map3k1* likely does not regulate viability. Together, *map3k1* inhibition led to formation of additional eyes at particular locations, with relatively normal composition and in an injury-independent manner.

### *map3k1* specifically regulates eye formation, not overall head patterning

In order to understand *map3k1*’s role in the process of eye formation and regeneration, we performed a series of comparisons between the *map3k1(RNAi)* phenotype and previously described ectopic eye phenotypes that cause either anterior (*notum* RNAi) or posterior patterning defects (*wnt11-6/wntA* RNAi and *ndk* RNAi) within the head that also impact brain:body regionalization. We first stained *map3k1(RNAi)* animals for *cintillo*^+^ cells, a population of antero- lateral neurons involved in mechano- or chemo-sensation and whose numbers scale proportionally to body size (30). While *ndk(RNAi)* animals underwent a posterior expansion and increased in average number of *cintillo^+^* cells per body area as expected, *map3k1(RNAi)* animals had no change in AP position and average number of *cintillo^+^* cells proportional to body area as compared to control animals (Fig 2A). Similarly, *ndk* RNAi caused posterior expansion of brain branches marked by *GluR^+^*, while *map3k1* inhibition did not appear to cause posterior brain expansion (Fig 2B). Additionally, *map3k1* RNAi did not cause any identifiable increases in *ChAT+* cholinergic neurons (Fig S2). These results suggest *map3k1* is unlikely to regulate eye production because of a role in global head patterning. To further examine this hypothesis, we tested markers of body regional identity to determine whether *map3k1* might have other roles in AP body axis formation.

**Fig 2.**
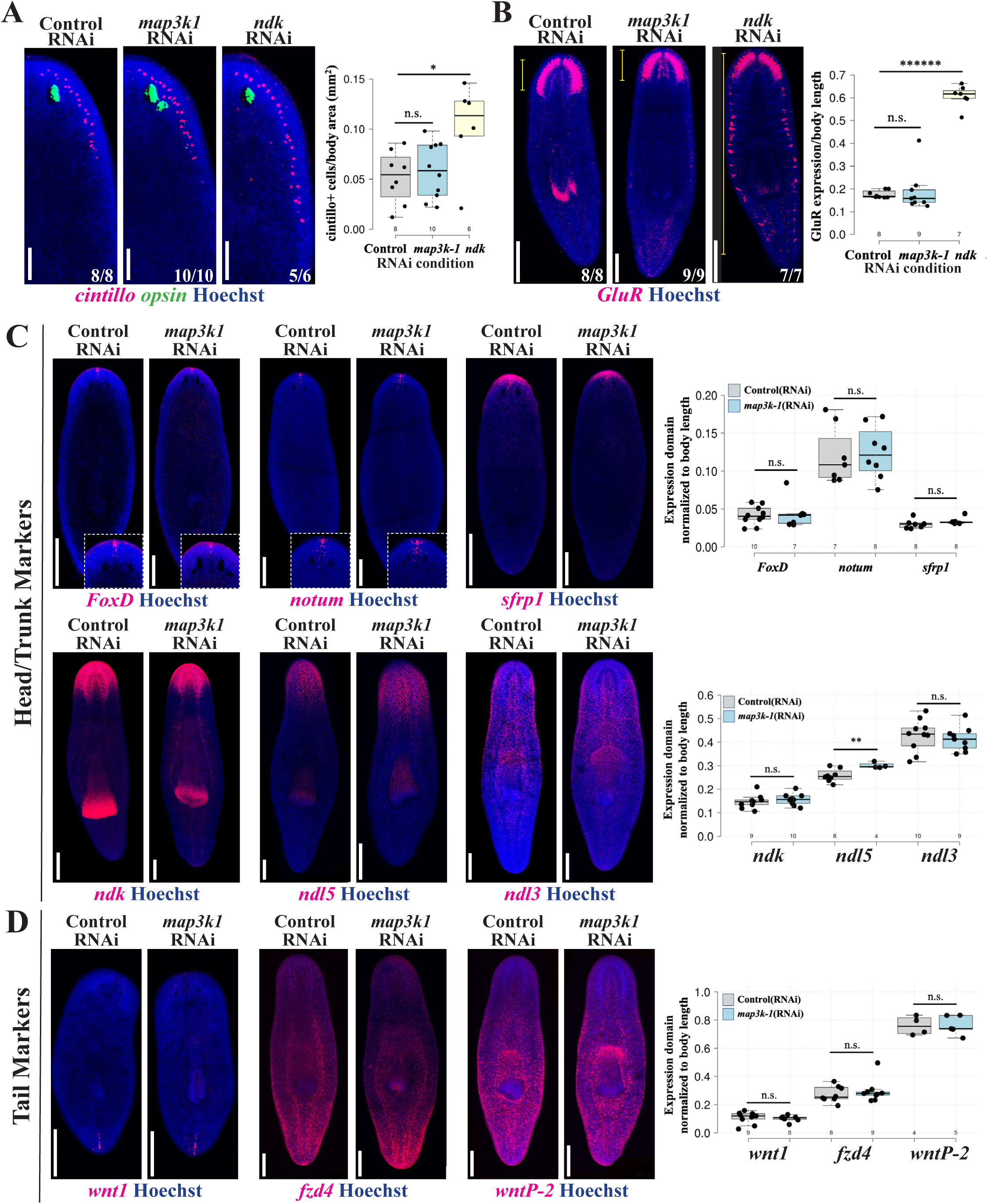
*map3k1* RNAi does not broadly affect brain size or body patterning. Homeostatic animals fixed after 3 or 4 weeks of RNAi were stained with markers to detect the effects of *map3k1* on brain and body patterning. (A) *ndk* RNAi caused an expected increase in the domain size (left images) and number (right quantifications) of brain-associated *cintillo* cells, while *map3k1* RNAi had no effect. Right, box plots overlayed with jittered scatterplots showing individual datapoints displaying the per-animal number of *cintillo^+^* cells normalized to body area in millimeters^2^ (mm^2^). *, p<0.05; n.s. represents p>0.05 as calculated by 2-tailed unpaired t-test; sample sizes for each condition are n≥6. Scale bars, 100μm. (B) Likewise, *map3k1* RNAi did not change the AP distribution of brain branches marked by *GluR,* while *ndk* RNAi resulted in ectopic brain branches forming throughout the body. Right, box plots showing the distance from the anterior tip of the animal to the most posterior *GluR* expression in the brain, relative to body length, for each of the corresponding RNAi conditions. ******, p<1E-7, n.s. represents p>0.05 as calculated by 2-tailed unpaired t-test; sample sizes for each condition are n≥7. Scale bars, 300μm. (C-D) Control and *map3k1(RNAi)* animals stained for position control gene (PCG) markers of (C) anterior and (D) posterior body patterning. Expression domains were either assessed after 14 days of regeneration following 2 weeks of dsRNA feeding and amputation of tails (*wnt1*), or assessed in uninjured animals treated with dsRNA for 3 weeks prior to fixation (all other probes). Right, plots showing quantification of expression domain sizes normalized to body length in Fiji/ImageJ. The sizes of *foxD*, *notum, sfrp1, ndk,* and *ndl5* expression domains were measured starting from the head tip to the posterior-most boundary of expression. *ndl3* occupies a position that does not reach the tip of the head, so the AP extent of this domain was measured instead. *wnt1, fzd4,* and *wntp-2* expression domain sizes were measured from the anterior-most expression to the tip of the tail. Each condition had a sample size of at least 5 animals. **, p<0.01; n.s., p>0.05 as calculated by 2-tailed unpaired t-test. Scale bars, 300μm. *map3k1* RNAi did not cause a measurable change to the expression domains for the majority of genes tested, and caused a small but statistically significant increase in the expression domain of *ndl5* expression.

*map3k1* RNAi did not affect AP expression domains for a majority of regional markers, including anterior/head markers *foxD, notum,* and *sfrp,* head/trunk markers *ndk* and *ndl3,* as well as tail markers *wnt1, fzd4,* and *wntP-2* (Fig 2C-D). *ndl-5,* marking the head region, was the only exception, because *map3k1(RNAi)* animals had a slightly expanded *ndl-5* domain compared to control animals. However, *ndl-5* is unlikely to have a strong role on eye patterning by itself, because does not itself, because prior work found that *ndl-5* RNAi did not cause patterning phenotypes (20). Together, we conclude *map3k1* likely does not control global AP axis identity or head regional identity, and instead acts at a novel control point that primarily regulates eye formation at a regulatory step distinct from previously identified eye regulatory factors.

### *map3k1* likely functions independently from Wnt signaling in eye regeneration

Our analysis of the *map3k1* RNAi phenotype suggested this gene likely controls a distinct step in eye formation compared to patterning signals, and we used double-RNAi analysis to further examine this possibility. In uninjured animals and regenerating head fragments, RNAi of the Wnt antagonist *notum* causes an anterior shift in head patterning to produce a second set of anteriorly positioned eyes (18). We reasoned that if *map3k1* functioned obligately downstream of *notum,* for example to relay signals downstream of *wntA,* then dual inhibition of both *notum* and *map3k1* might tend to produce only the ectopic posterior eyes phenotype from *map3k1* RNAi. We inhibited *notum* and *map3k1* together under homeostatic conditions, and then compared this outcome to single-gene inhibitions spiked with competing control dsRNA. Under these conditions, *notum(RNAi)* animals all formed anterior ectopic eyes (13/13 animals, 100%) and *map3k1(RNAi)* animals formed ectopic posterior ectopic eyes (14/14 animals, 100%). By contrast, the majority of *notum;map3k1(RNAi)* animals displayed a synthetic phenotype in which both ectopic anterior and posterior eyes formed (14/15 animals, 93%). These outcomes argue that *map3k1* likely controls eyes formation at a step independent from notum/Wnt signaling and consequently head patterning (Fig 3).

**Fig 3.**
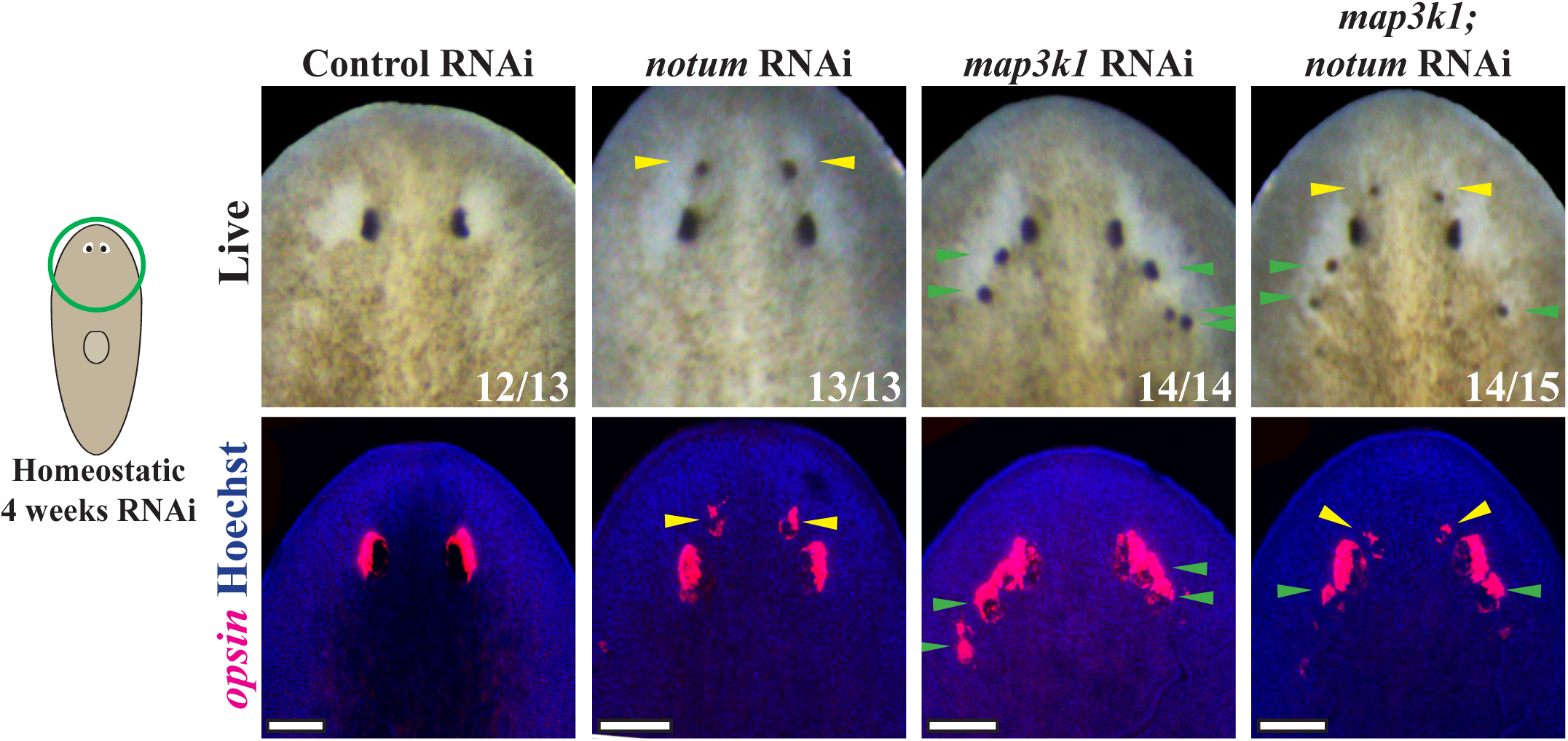
*map3k1* likely regulates eye formation independent of Wnt signaling. Double RNAi experiments were conducted to test potential interactions between *notum* and *map3k1* genes whose individual inhibition causes spatially distinct ectopic eye phenotypes. Animals were fed dsRNA every 2-3 days for 4 weeks before live scoring (top panels) and fixation to detect *opsin^+^* eye cells (bottom panels). As expected, *notum(RNAi)* animals formed anterior ectopic eyes (13/13), while *map3k1(RNAi)* animals formed posterior ectopic eyes (14/14). However, nearly all *map3k1;notum(RNAi)* animals formed a synthetic phenotype in which both anterior and posterior ectopic eyes formed (14/15). Therefore, it is likely that *map3k1* and Wnt pathways regulate distinct processes in eye formation. Yellow arrows are used to highlight anterior ectopic eyes while green arrows are used to highlight posterior ectopic eyes. Single-RNAi conditions involved combining control dsRNA with an equal amount of experimental dsRNA so that all treatments received the same total amount of dsRNA. Scale bars, 100μm.

In normal uninjured animals, eye removal by resection results in regeneration of a new eye at precisely the original location. By contrast, removal of eyes in animals inhibited for patterning factors *wntA, fzd5/8-4, src-1* or *notum,* eyes only regenerate at the location of the newly formed ectopic eyes, and not the pre-existing eyes, despite being able to be maintained for long periods of time homeostatically (18, 22). In principle, *map3k1* could control patterning within the head region but in a manner specific to eye placement and not brain patterning. To test this possible model, we measured the outcomes of original and ectopic eye removal in *map3k1(RNAi)* animals. In homeostatic *map3k1(RNAi)* animals, removal of the original eyes resulted in eye regeneration at that location in 100% of animals (Fig S3). Similarly, removal of ectopic eyes from these animals resulted in regeneration in these same locations in 50% of animals. Therefore, in these experiments, *map3k1* inhibition did not shift the target location for eye regeneration away from the original eyes. Instead, eye regeneration became permissible in an expanded domain. These outcomes further support a model in which *map3k1* likely acts independently of Wnt/*notum* signaling and suggests a role distinct from known patterning factors.

### *map3k1* is expressed broadly throughout the body, including in eye cells

We then investigated the expression of *map3k1* transcripts in the body, with particular interest with whether this factor is expressed in differentiated eye cells, undifferentiated eye progenitors, or other cell types. By analysis of published single-cell RNAseq data, *map3k1* expression was broad and present in many cell types, including muscle, gut, neurons, and protonephridia (Fig S4A) (31). FISH using a *map3k1* riboprobe displayed broad staining across most regions of the animals, putatively in locations around the brain, flame cells, and also near the eyes, based on anatomical location (Fig S4B). We used double-FISH with *opsin* and *ovo* to verify whether *map3k1* is expressed in differentiated eye cells and/or eye progenitors. *map3k1* FISH signal was detected throughout the *opsin^+^* cells in all animals (4/4, 100%) (Fig S4C) and also weakly within *ovo^+^* progenitors (Fig S4D, yellow arrows). We also noticed some nearby ovo^-^ cells of unknown cell type that had higher levels of *map3k1,* which could either represent cells of unknown function in eye regeneration or unrelated cells (Fig S4D, yellow arrows). These results are consistent with *map3k1* acting either within mature eye cells, migratory eye progenitors, or could indirectly regulate eye formation through action in an alternative cell type.

### *map3k1* RNAi increases numbers of differentiated eye cells

We reasoned that rather than controlling eye and head patterning, *map3k1* might instead limit the extent of eye cell differentiation. Prior work indicated that the emergence of ectopic organs in planarians does not necessarily correspond to increased cell type production. For example, *notum* RNAi pattern transformations can result in the formation of extra eyes without increasing numbers of total eye cells throughout the animal (18), suggesting pattern alteration can proceed in a way that spatially re-partitions a defined number of differentiating cells. To investigate *map3k1’s* influence on eye cell abundance, we quantified the total number of *opsin^+^*differentiated eye cells in control versus *map3k1(RNAi)* homeostatic animals. To do this, we sampled regularly-spaced confocal slices from z-stacks of the animal eyes, detected number of eye cells per slice using 2D segmentation of nuclei followed by measuring overlap with *opsin+* FISH signal after applying thresholding, and summed *opsin+* nuclei across slices sampled for each animal (See methods, Fig 4A). This analysis revealed that *map3k1(RNAi)* animals contained significantly more differentiated *opsin^+^* cells compared to control animals (Fig 4B-C) and were also distributed in a broader spatial range compared to control animals. The detection of an increase in *opsin*^+^ cell numbers after *map3k1* RNAi was also robust to variation of the exact threshold and inter-slice distance chosen (see Methods). To examine the possibility of any systematic differences in segmentation quality across the two treatments, we calculated a Jaccard Similarity Index (JSI, also known as the “Intersection over Union” metric) on randomly selected animals and z-slices, then compared these metrics between control and *map3k1(RNAi)* conditions. We found that the mean JSIs for control and *map3k1(RNAi)* samples were 0.55 and 0.49, respectively, and did not differ from each other as measured by a 2-tailed unpaired t-test (p=0.23, Table S2). Therefore, the differences in estimated cell numbers were not likely due to any systematic differences in the segmentation and counting efficiency across the treatments. We conclude that *map3k1* RNAi causes an increase in total *opsin+* eye cell numbers per animal, in addition to causing the formation of ectopic eyes. Together, these observations suggest that *map3k1* normally limits both the location and extent of eye cell differentiation.

**Fig 4.**
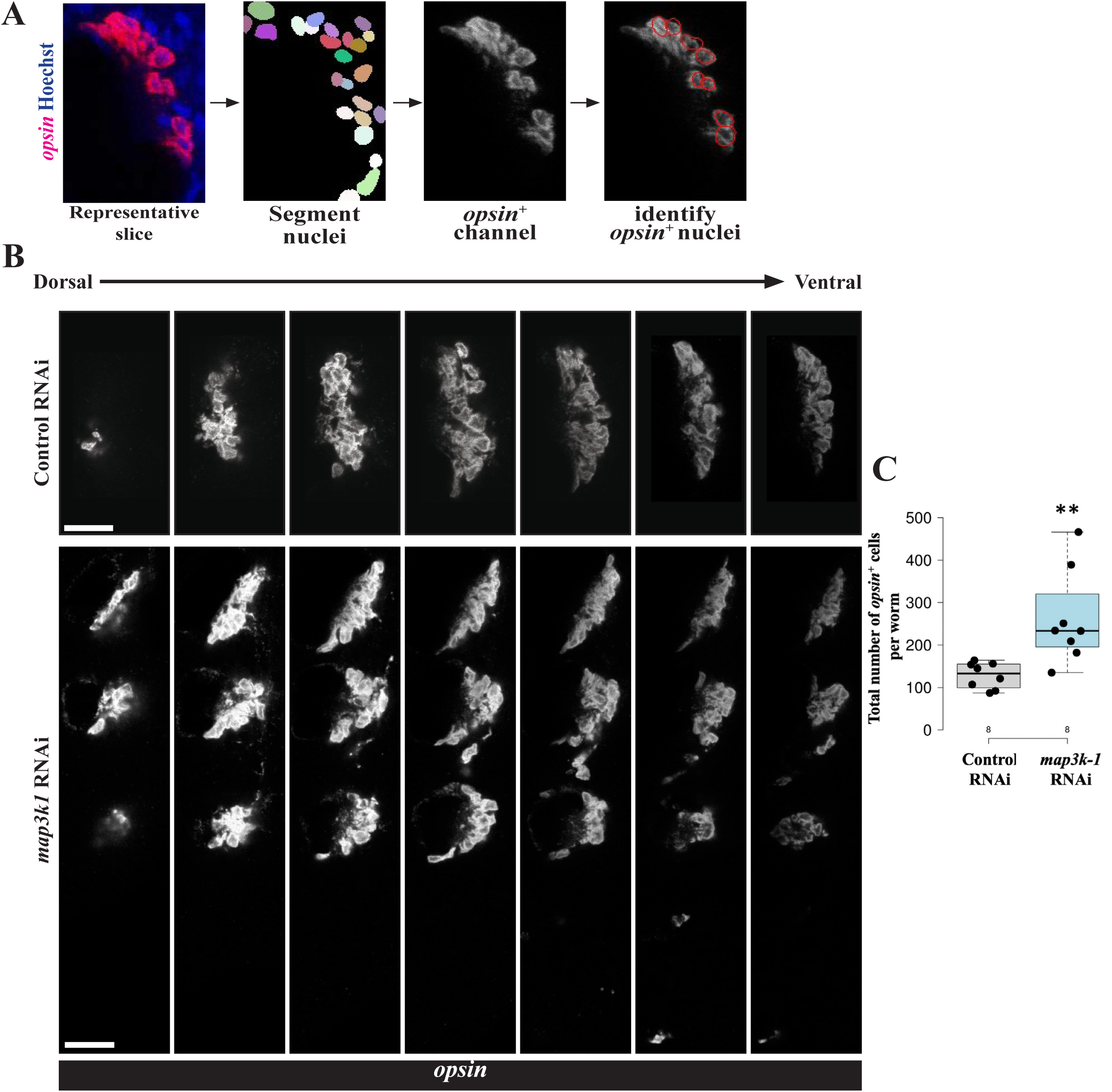
*map3k1* inhibition causes an increase in numbers of differentiated eye cells. Uninjured control and *map3k1(RNAi)* animals fixed after 3 weeks of RNAi and stained with *opsin* riboprobe to quantify numbers of eye cells in each condition. Eye cells are present in close association with each other, necessitating an image analysis workflow for their quantification. Z-stacks capturing all eye cells were obtained through confocal imaging, then slices 5-microns apart were selected to represent each stack for 2D segmentation using Stardist, followed by assignment of nuclei as *opsin*^+^ using a global threshold and summing number of positive cells across the selected stacks for each animal. (A) Example of an image slice after nuclei segmentation and assignment of *opsin^+^*nuclei (red overlay, right) of how *opsin^+^* cells were counted for one z-stack. (B) Scaled images of example z-stacks from control and *map3k1(RNAi)* animals showing that *map3k1* inhibition resulted in an expansion of eye regions. (C) Total number of *opsin*^+^ cells counted in control versus *map3k1(RNAi)* animals. *map3k1* inhibition caused an increase to the number of measured eye cells. Plots show data points overlaid with boxplots. **p=0.01 by 2-tailed unpaired t-test. n=8 animals. Scale bars, 25μm.

### *map3k1(RNAi)* ectopic eyes form in regions normally occupied by eye progenitors

We hypothesized that *map3k1* might control the terminal differentiation of eye progenitors and reasoned that the locations of ectopic eye formation might provide additional information useful for distinguishing this role from the patterning systems regulating eye positioning. We noted that *map3k1(RNAi)* ectopic eyes seemed to emerge stochastically in a particular region of the body laterally and posteriorly to the normal eye location. By contrast, extra eyes forming after inhibition of *wntA, ndk,* or *notum* often appear symmetrically and in specific locations (18, 21). To examine these distributions, we systematically quantified eye position in images of live animals following homeostatic RNAi of *map3k1, wntA, notum,* and *ndk*. Because planarians lack a fixed size and readily grow or de-grow while maintaining proportionality, we normalized the AP and ML position of each ectopic eye with respect to the original eye locations in each animal (see Methods). This analysis confirmed that *map3k1(RNAi)* ectopic eyes generally form across a broad region located laterally and posterior to the original eyes, whereas the majority of *wntA(RNAi)* and *ndk(RNAi)* ectopic eyes form in a more concentrated region located directly posterior to the original eyes (Fig 5A-B). This analysis provides further confirmation that *map3k1* likely controls a distinct step in eye formation compared to *wntA* and *ndk*.

**Fig 5.**
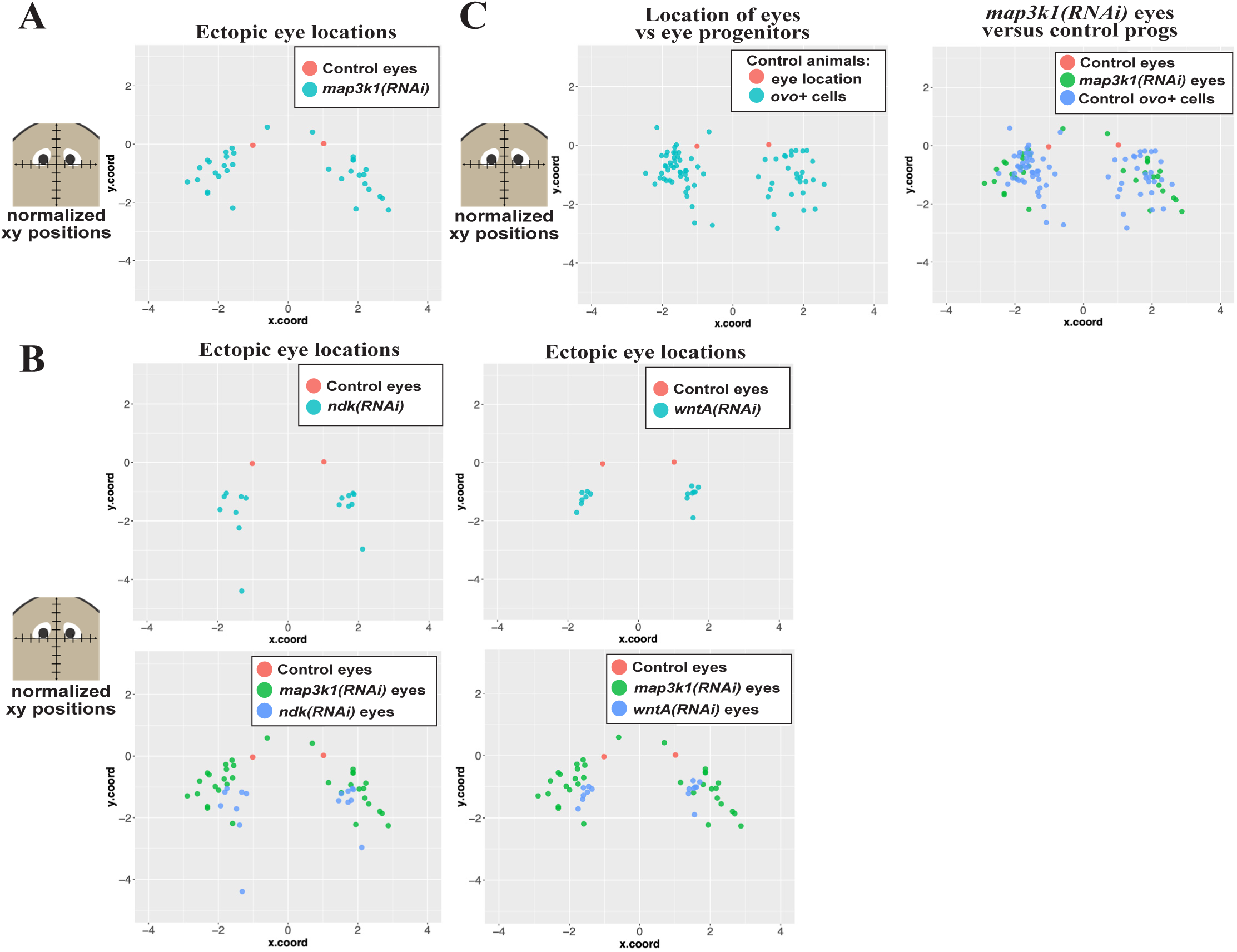
*map3k1(RNAi)* ectopic eyes form in a posterior and lateral region within the domain of normal *ovo^+^* migratory cells. Homeostasis animals were treated with the indicated dsRNA for 3 weeks (control RNAi, *wntA* RNAi) or 6 weeks (control RNAi, *map3k1* RNAi, *ndk* RNAi) followed by live imaging to detect the location of eyes or fixing and staining to detect the location of migratory *ovo^+^* cells (eye progenitors) in unfed uninjured animals. Planarians lack a fixed size, so in order to make comparisons across treatments, locations of ectopic eyes and eye progenitors in each image were defined and then registered and normalized to the location of the original eyes in order to create a common coordinate system (See methods). These data were plotted as indicated (A-C) in which the AP (y.coord) and ML (x.coord) axes are plotted with units equal to one-half of the inter-eye distance as measured either between the normal eyes of control animals or between the original eyes in *ndk(RNAi)*, *wntA(RNAi)*, and *map3k1(RNAi)* conditions. (A) Scatterplot of control eyes (red dots, from 5 animals) versus *map3k1*(RNAi) ectopic eyes (light blue dots, from 8 animals) shows that *map3k1* inhibition caused formation of ectopic eyes in a distribution located laterally and posteriorly compared to control eye locations. However, rare *map3k1(RNAi)* ectopic eyes were also identified slightly anterior to the original eyes (2 blue dots with y.coord>0). (B, top) Plots of ectopic eye locations in *ndk(RNAi)* or *wntA(RNAi)* animals (light blue dots) with respect to control eyes (red dots). (B, bottom) Graphs comparing locations of ectopic eyes in *map3k1(RNAi)* (green dots) versus either *ndk(RNAi)* animals or *wntA(RNAi)* animals (blue dots). Both *ndk* RNAi and *wntA* RNAi caused a tighter distribution of ectopic eyes that were located more directly posterior to the original eyes compared to the broader distribution of *map3k1(RNAi)* ectopic eyes located more laterally. (C, left) Locations of migratory *ovo^+^*eye progenitors from control uninjured animals (red dots) were compared to *ovo^+^* mature eyes (light blue dots). (C, right) Locations of *ovo^+^* eye progenitors (blue dots) were compared to locations of ectopic eyes from *map3k1(RNAi)* animals (green dots). *map3k1* inhibition caused formation of ectopic eyes in a set of locations overlapping with the location of the eye progenitors from control animals.

The position of *map3k1(RNAi)* ectopic eyes were instead reminiscent of the distribution of *ovo^+^* eye progenitor “trails” leading to the eye that are most evident 5-8 days after decapitation, during the late stages of head regeneration (6). We hypothesized that if *map3k1* limits the terminal differentiation of migratory progenitors without influencing patterning, then ectopic eyes in *map3k1(RNAi)* animals might be most likely to form within regions that ordinarily contain *ovo^+^* cells. To make a systematic comparison, we employed a similar positional mapping strategy used above, to measure the location of *ovo+* cells in homeostatic control animals, then plotted this distribution in comparison to the position of *map3k1(RNAi)* eyes (Fig 5C). *map3k1(RNAi)* ectopic eyes were located within a subset of the region occupied by *ovo*^+^ cells in control animals (Fig 5C). In the process of undertaking this analysis, we noticed a few very rare occurrences in which ectopic eyes from *map3k1(RNAi)* animals formed anterior to the original eyes (2/8 animals had 1 anterior eye each in this experiment, Fig 5A). These observations, while encompassing only a small fraction of eyes surveyed, provide additional evidence that *map3k1* likely does not exclusively control the process of anterior versus posterior eye placement. Similarly, the majority of *ovo^+^* cells were posterior to the existing eyes, as reported previously, but rare *ovo^+^* cells could also be detected anterior to the eyes. This observation is consistent with a model in which *ovo^+^* eye progenitor specification normally takes place over a broad anterior region, including to some extent anterior to the eyes, but receive signals which hone progenitors to migrate and incorporate into eyes for cell replacement and growth (6, 25). Furthermore, the distribution of *map3k1(RNAi)* ectopic eyes is consistent with a model in which *map3k1* normally maintains *ovo^+^* migratory progenitors in an undifferentiated state prior to arrival at the precise location of terminal differentiation.

### *map3k1* suppresses differentiation of migratory *ovo*^+^ eye progenitors

Based on these findings, we hypothesized *map3k1* might regulate terminal differentiation rather than the initial specification of *ovo^+^* cells. To test this possibility, we examined the number of *ovo^+^* cells across different stages of head regeneration, beginning with *ovo*^+^ cell specification from neoblasts within 2-3 days of injury to their subsequent localization into posterolateral trails that support the differentiation of new eye cells within the blastema. In *map3k1(RNAi)* trunk fragments, a single set of seemingly normal eyes formed by 3-7 days after amputation, followed by formation of ectopic eyes after 2 weeks of regeneration (Fig 6). We used a FISH experimental strategy to quantify numbers of progenitors during this process. Double-FISH simultaneously detecting *ovo* along with a mixture of *opsin* and *tyrosinase* riboprobes allowed distinguishing progenitors from mature eye cells. We then compared the number of *ovo^+^* progenitor cells in control versus *map3k1(RNAi)* trunk fragments across a time series of regeneration (Fig 6). The number of undifferentiated *ovo^+^* eye progenitors were unchanged between *map3k1(RNAi)* and control animals at days 0, 2, 5, and 8. However, *map3k1* RNAi caused a decrease to the number of *ovo^+^* eye progenitors at 14 days post-amputation, at a time coinciding with the emergence of ectopic eyes in these animals. Taken together, *map3k1* inhibition can lead to decreases in progenitors and also increases in terminally differentiated cells. These results indicate that *map3k1* likely does not function in the specification of neoblasts into migratory eye progenitors, but instead regulates the ability for migratory eye progenitors to remain undifferentiated in regeneration.

**Fig 6.**
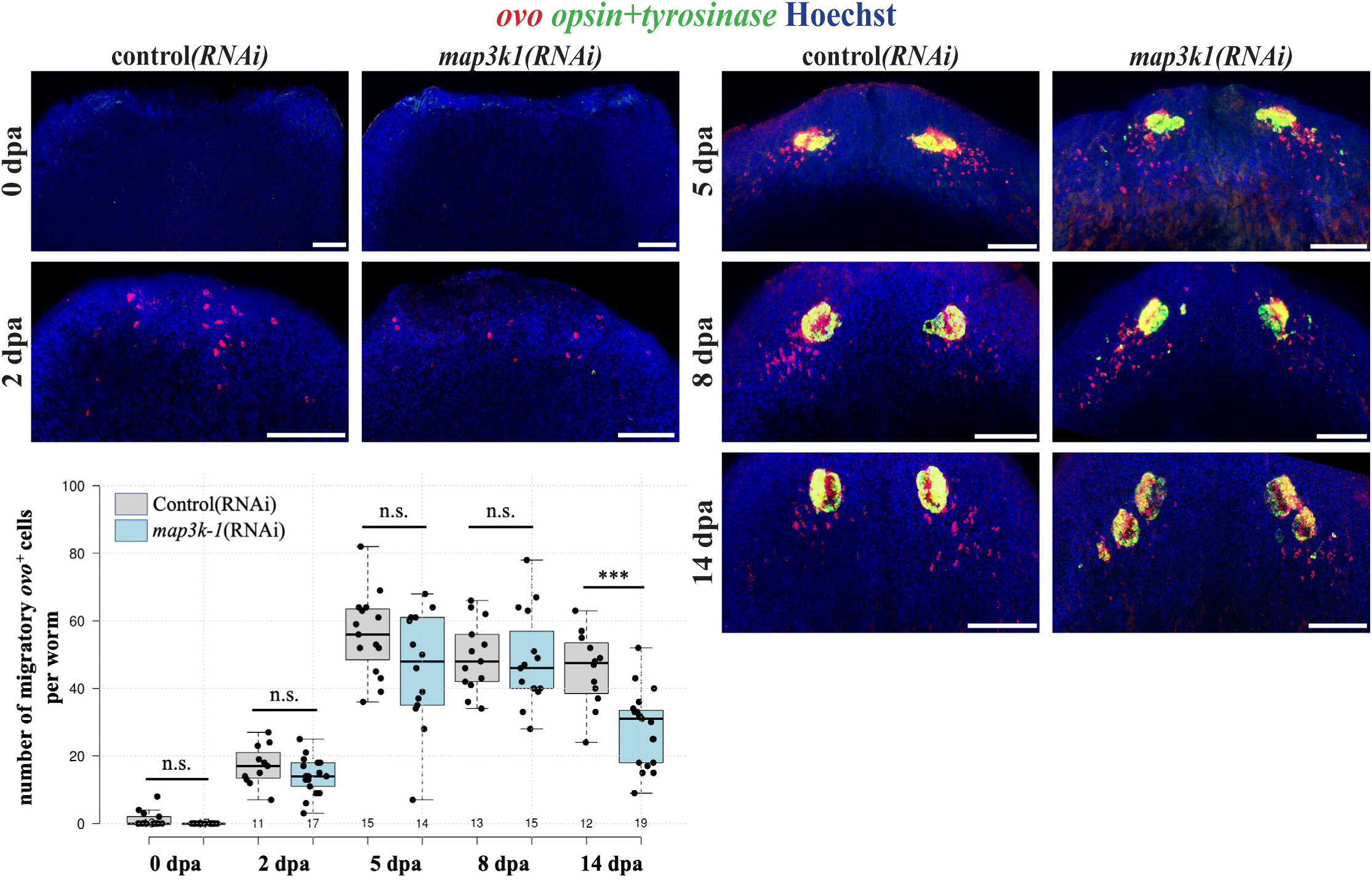
*map3k1* inhibition reduces number of undifferentiated *ovo^+^* eye progenitors during formation of ectopic eyes. Animals were treated with control and *map3k1* dsRNA for 2 or 3 weeks prior to amputation of heads, and the resulting regenerating trunk fragments were fixed in a time series followed by staining with an *ovo* riboprobe to detect migratory eye progenitors and simultaneously with mixture of *opsin* and *tyrosinase* riboprobes to detect mature eye cells. Numbers of *ovo^+^* migratory progenitors were quantified for each timepoint and condition using maximum projections. Data for 0dpa and 2dpa were aggregated from two experiments. Data for 5dpa, 8dpa, and 14dpa were aggregated from two different experiments, each showing decline in *ovo^+^* eye progenitor numbers at 14dpa. The number of undifferentiated *ovo*^+^ eye progenitors did not change significantly in *map3k1(RNAi)* worms at 0, 2, 5, and 8dpa. At 14dpa, a time when ectopic eyes began to emerge, *map3k1(RNAi)* worms showed a decrease in *ovo^+^* eye progenitors. Plot show data points from individual animals overlaid with boxplots. ***p<0.001, n.s. represents p>0.05 by a 2-tailed unpaired t-test. Scale bars, 100μm.

### *map3k1* likely signals through *map2k-4* and *map2k-7* to suppress eye formation

We used the molecular identity of *map3k1* to inform targeted secondary RNAi screens in order to determine a candidate pathway of action for this gene in eye formation. In a typical MAP Kinase pathway, MAP3Ks signal to MAP2Ks, which then activate MAPK in order to regulate downstream responses. To identify the *map3k1* pathway relevant for eye formation in planaria, we first identified all 8 MAP2K genes then screened these individually by RNAi to determine whether any would phenocopy the ectopic eye effect. However, knockdown of planarian MAP2K genes individually did not produce any eye phenotypes (Fig 7A), suggesting a potential redundancy in their uses. We then inhibited combinations of genes and found that *map2k4;map2k7(RNAi)* animals formed ectopic eyes in regenerating head fragments (4/10, 40%) similar in appearance, though at a lower penetrance, to the effects of *map3k1* RNAi (Fig 7B). MAP2K4 and MAP2K7 are known to transduce mammalian MAP3K1 signals, and frequently do so by regulating p38 and/or JNK downstream MAPKs (32). We therefore inhibited the two planarian *p38* genes in combination, as well as *jnk*, by administering 6 dsRNA feedings over 2 weeks followed by 14 days of regeneration. Under these conditions, inhibition of *jnk* or both homologs of *p38* caused ectopic eye phenotypes similar to *map3k1* inhibition, though again at a low penetrance (Fig 7C). The reduced penetrance from inhibitions of downstream MAP2K and MAPK factors, compared to *map3k1* inhibition, could indicate further uncharacterized redundancy, or perhaps inefficient protein knockdown from RNAi. These factors likely also have independent uses and inputs in planarians, as previously argued from other studies using alternate dsRNA dosing schedules and methods to perturb JNK and p38 signaling (33–36). However, our results argue that *map3k1* likely signals via MAP2K4/MAP2K7 and JNK/p38 in order to suppress differentiation of planarian eye cells.

**Fig 7.**
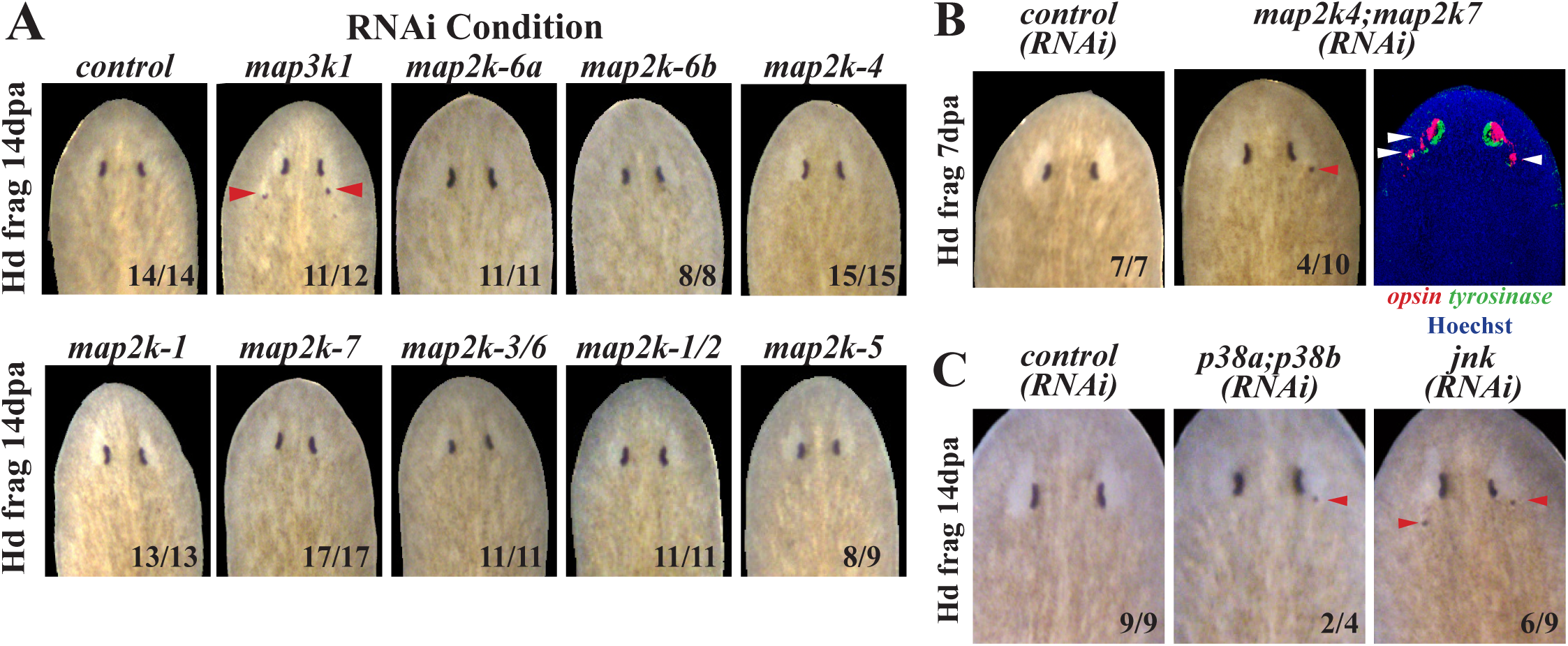
*map3k1* likely controls eye progenitor differentiation through a MAP2K4/7-JNK/p38 pathway. Members of MAP2K and MAPK gene families were inhibited in order to identify signals downstream of *map3k1* regulating eye formation. Animals were treated with indicated dsRNAs for 2 weeks (A-B) or 3 weeks (C) prior to amputation of tails and allowed to regenerate for 7 days (B) or 14 days (A, C) followed by live imaging. (A) While *map3k1(RNAi)* animals developed ectopic eyes as expected (red arrows), single gene inhibitions of *map2k* genes did not cause formation of ectopic eyes. (B) *map2k4;map2k7(RNAi)* animals were observed to develop posterior ectopic eyes (red arrow) through live imaging. Double FISH detecting *opsin^+^* photoreceptor neurons and *tyrosinase^+^* pigment cup cells showed that the ectopic eyes contained both *opsin^+^* and *tyrosinase^+^* cells (white arrows). (C) Inhibition of both homologs of the planarian *p38,* or *jnk*, caused ectopic eyes to form in regenerating head fragments. Scorings indicate the number of animals displaying the phenotype depicted in each panel.

### *map3k1* levels may control the locations of terminal differentiation

In planarians, undifferentiated eye progenitors appear to migrate over potentially large distances to reach the eye. In uninjured animals, *ovo*^+^ cells can be detected sporadically within the anterior half of planarians, and these animals scale between 2-20mm in length. The “trail” region in which *ovo^+^* cells are more concentrated, and presumably represent late phases of migration into the eyes, would occupy about 10% of the AP axis for a range of approximately 1∼2 mm. Given the analysis supporting a role of *map3k1* in maintaining the undifferentiated state during a potentially lengthy migration, we wondered whether this gene was equally important at all stages during cell transit. In principle, an immediate switch to terminal differentiation across all eye progenitors should result in ectopic eyes forming uniformly across the field of migrating cells. However, we noted that the *map3k1(RNAi)* ectopic eyes seemed to form in a progressively posterior manner during the RNAi treatment, suggestive of some progressive requirement for this gene as eye progenitors reach their target. Based on these results, we hypothesized that progenitors closer to the eyes versus further from the eyes might have different requirements for *map3k1* levels. To test this hypothesis, we designed an experiment to investigate the outcome of reduced *map3k1* silencing on the ectopic eye phenotype by comparing homeostatic animals treated with varying amounts of *map3k1* dsRNA along with competing control dsRNA, analogous to a genetic allelic series. If all eye progenitors across the head were equally sensitive to *map3k1* inhibition, we expected that weakened silencing would result in the same AP distribution of ectopic eyes but at a lower penetrance. By contrast, compared to the strongest *map3k1* dsRNA treatment (“100%”), treatment with the reduced dosage of *map3k1* dsRNA (“50%”) caused formation of fewer ectopic eyes (Fig 8A) and reduction in the posterior extent of ectopic eye formation (Fig 8B-C). Therefore, it seems that eye progenitors in different AP regions of the head are differentially susceptible to *map3k1* silencing. Because RNAi silencing in planarians is not known to have an AP bias in general (37), these results suggests that relatively less *map3k1* silencing is required to cause terminal differentiation of eye progenitors located at more anterior positions closer to the target destination as compared with more posterior eye progenitors. A possible explanation for this effect could be that in normal animals, signaling through *map3k1* attenuates as progenitors reach their target, leading to terminal differentiation coinciding with the target location of the mature eyes. These results suggest that *map3k1* participates helps to coordinate the spatial locations of terminal eye cell differentiation.

**Fig 8.**
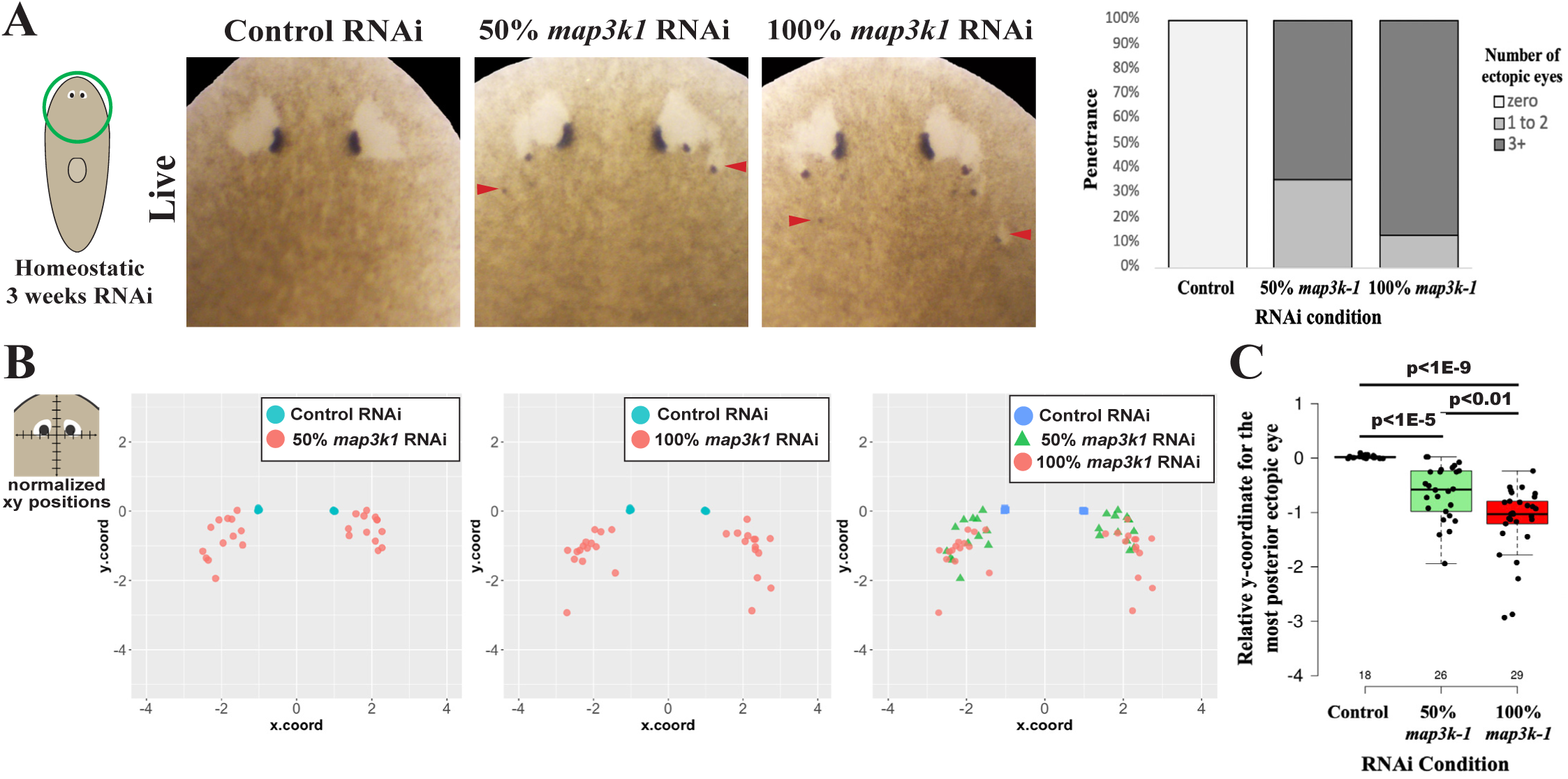
Increasing the dose of *map3k1* dsRNA causes ectopic eyes to form more posteriorly. Uninjured animals were treated with 3 weeks of either control dsRNA, a mixture of 50% *map3k1* dsRNA combined with an equal amount of competing control dsRNA, or 100% *map3k1* dsRNA in order to test how different levels of *map3k1* inhibition affect the AP locations of ectopic eye formation. (A) Images of live animals after the indicated treatments (left panels), and scoring of the fraction of total animals (% penetrance) obtaining zero, versus 1-2, versus 3 or more ectopic eyes, plotted in a stacked bar graph (right panels). Animals receiving higher doses of *map3k1* dsRNA formed a greater number of ectopic eyes. (B) To determine whether the spatial distribution of ectopic eyes was dependent on the dose of *map3k1* dsRNA, relative AP and ML positions of the posterior-most ectopic eye in each animal were quantified as in Figure 5 and visualized in scatterplots (see Methods). This procedure normalized the AP (y.coord) and ML (x.coord) positions of the posterior-most ectopic eye in each *map3k1(RNAi)* animal compared to the location of the original eyes, with units corresponding to one-half of the inter-eye distance in control*(RNAi)* animals or of the original eyes in *map3k1(RNAi)* animals. (B, left panels) Plots of the most posterior ectopic eye locations in 50% *map3k1(RNAi)* or 100% *map3k1(RNAi)* animals (red dots) with respect to control eyes (light blue dots). (B, right panel) Scatterplots comparing locations of ectopic eyes in 50% *map3k1(RNAi)* (green triangle) versus 100% *map3k1(RNAi)* animals (red dot). (C) Boxplot shows the AP locations (y.coord) of the posterior-most eye in each animal, with individual datapoints overlayed as a jittered scatterplot. p-values as calculated by an unpaired 2-tailed t-test are indicated. Higher doses of *map3k1* dsRNA caused ectopic eyes to form in more posterior locations. Sample size, n≥11 animals for each condition.

## Discussion

Our analysis identifies a critical and novel role for *map3k1* in controlling the site of eye progenitor differentiation during eye regeneration in planarians (Fig 9). Several lines of evidence suggest *map3k1* controls a distinct step in eye regeneration compared to other known patterning factors. *map3k1* silencing caused eye-specific effects rather than also affecting brain:body patterning, *map3k1* interacted synthetically with *notum,* the locations of ectopic eyes in *map3k1(RNAi)* differed from those in *wntA(RNAi)* and *ndk(RNAi)* animals, and *map3k1* RNAi did not cause the target location for eye regeneration did not shift away from the location of the original eyes. Additionally, we found that *map3k1* inhibition can lead to increases in numbers of terminally differentiated eye cells, and decreases to eye progenitor numbers in regeneration. These results together suggest that *map3k1* signaling regulates a novel control point involved in preventing terminal differentiation during the migration of eye progenitors to their target destinations.

**Figure 9.**
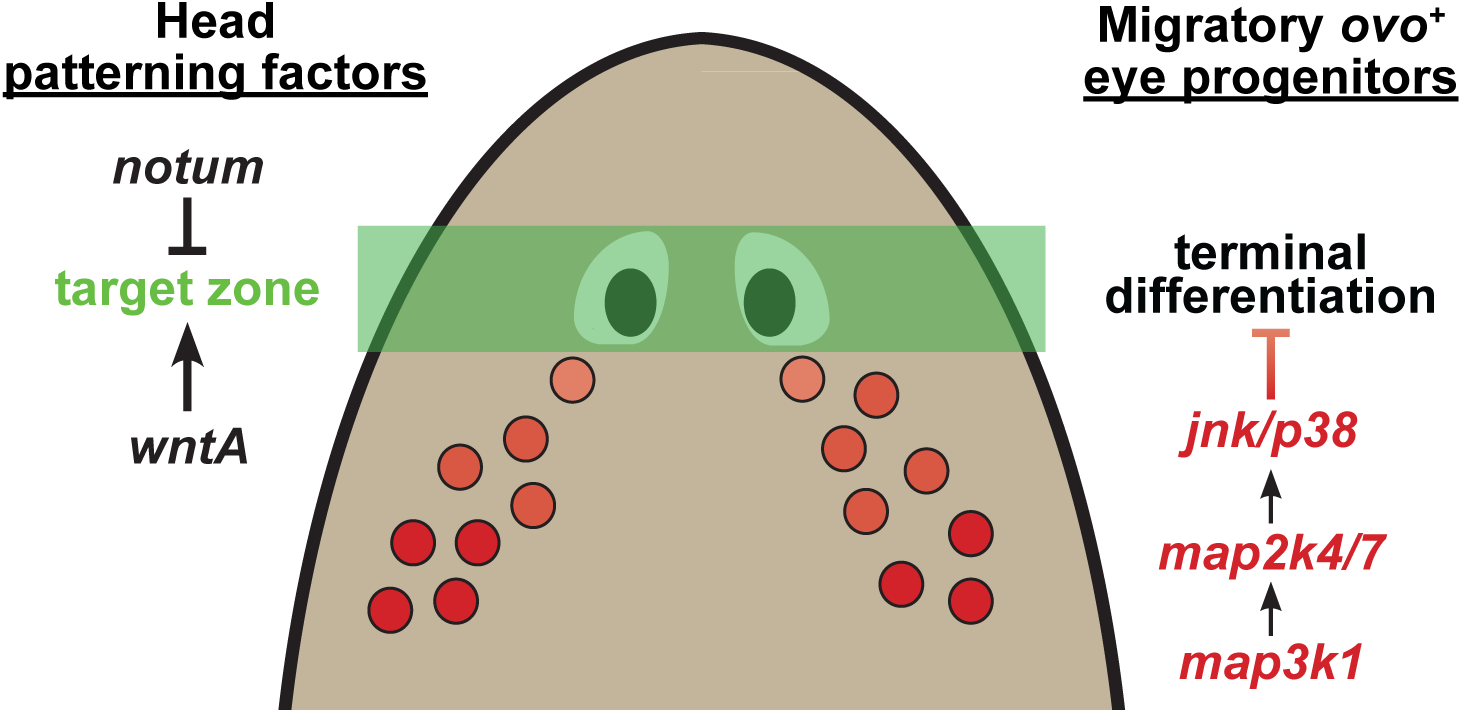
*map3k1* suppresses terminal differentiation of migratory eye progenitors. Model of the role of *map3k1* in eye regulation. Head patterning factors such as *notum* and *wntA* regulate a target zone for eye placement in regeneration, while *map3k1* suppresses terminal differentiation of migratory eye progenitors by activating Map2k4/Map2k7-JNK/p38 downstream signals. dsRNA dosage experiments indicate that progenitors located further away from the eyes are less susceptible to the effects of *map3k1* inhibition, suggesting that *map3k1*-dependent processes may normally decline in activity as eye progenitors reach their destination.

Map3k1 proteins in other organisms are known to be involved in control of multiple cellular outputs, including promoting cell survival and enabling migration. Mammalian *Map3k1* was originally identified as a spontaneous mutant in 1966 called lidgap-Gates (lg^Ga^) which displayed an eyes-open-at-birth (EOB) phenotype caused by a failure or delay in epithelial migration of the eyelid epidermis. lg^Ga^ was subsequently mapped and cloned as a deletion of Map3k1 (38, 39). Targeted mutations found that mice lacking the full-length *map3k1* gene (*Map3k1*^-/-^) and also mice lacking the *map3k1* kinase domain (*Map3k1^τιKD/τιKD^*) both display an EOB phenotype due to improper cell-cell interactions and failed migration of eyelid epithelial cells (27, 40, 41). Eyelid closure involves a collective migration of epidermal cells and inputs from many pathways, including EGF, FGF, Wnt, TGF-beta, and JNK signaling (41–45). However, the due to the complexity of signals involved in this process, the precise signals impinging on Map3k1 are not yet fully understood. Interestingly, Map3k1 genes are not present in *Drosophila melanogaster* or *Caenorhabditis elegans* genomes, suggesting planarians could be a useful system for understanding the roles this pathway plays in differentiation.

What are the developmental mechanisms underlying *map3k1*’s role in differentiation? Prior work found that in the planarian *Dugesia japonica*, *map3k1 (Djmekk1)* was required for correct positioning of trunk tissues during regeneration (46). In principle, these roles could be related to the control of terminal differentiation our study identifies, for example, if distinct RNAi dosing strategies reveal a broader role of *map3k1* in migratory progenitor maintenance relevant to tissues beyond the eyes. It is likely that the *Schmidtea mediterranea map3k1* participates in signaling beyond eye formation due to the fact this gene is broadly expressed. Thus, identifying the relevant upstream and downstream factors acting through *map3k1* will be important for resolving the developmental mechanisms by which this factor controls eye regeneration. Previous research identified that Map3k factors can be activated through a wide variety of signals, including EGF, FGF, and also cold shock, microtubule disruption, or LPA treatment, and pathways downstream of *map3k1* can include MAP2K4 and MAP2K7 signaling to p38 or JNK (40, 41, 47, 48). While we not able to identify signals upstream of *map3k1,* we found that inhibition of *map2k4* and *map2k7*, *jnk*, or *p38* could cause a low-penetrance phenocopying of the *map3k1* RNAi ectopic eye phenotype. Prior analysis of JNK or p38 signaling in planarians has also revealed their participation in several aspects of regenerative growth, including cell-cycle control, differentiation, and regulation of apoptosis (34–36, 49, 50). While we did not observe similar effects, our analysis reveals the potential existence of a Map3k1-Map2k4/Map2k7-JNK/p38 signaling pathway in planarians with a prominent role in controlling eye cell terminal differentiation.

Some aspects of the *map3k1(RNAi)* phenotype suggest complexity in the control of terminal differentiation used in regeneration. For example, in our experiments, knockdown of *map3k1* did not result in complete elimination of all migratory eye progenitors through their terminal differentiation, as we were able to observe *ovo^+^* undifferentiated cells which were still present in *map3k1(RNAi)* animals. In addition, our experiments to investigate the effects of weaker versus stronger inhibition of *map3k1* revealed that progenitors closer to their destination were more likely to differentiate into ectopic eyes. Together, this suggests that some progenitor cells are more susceptible to the effects of *map3k1* inhibition than others. However, whether these results arise because of a spatial gradient of *map3k1* activity, or some other spatially related effect, remains to be determined. We cannot rule out the possibility that these observations arise from undetected ways that RNAi effects spread differentially across the planarian body. However, phenotypes that simultaneously affect anterior and posterior regions have been described previously (37), suggesting this explanation is unlikely. Instead, we suggest that either Map3k1 signaling itself, or alternatively some Map3k1-dependent process, could attenuate as progenitors in normal animals reach the eye, which would coordinate the process of terminal differentiation to occur at the correct location. Interestingly, gradients of mammalian Map3k1 gene activity have been proposed to operate in eyelid formation, on the basis of genetic enhancement observed from interaction of heterozygous alleles of Map3k1 and JNK in producing eyes-open-at-birth phenotypes (51). Developmental activity gradients of mammalian Map3k1 protein function have not been described, but the planarian eye regeneration paradigm could be a useful system to investigate the mechanisms that enable the appropriate timing and location of progenitor differentiation. Another possibility is that *map3k1* signals are critical for a subset of eye progenitors primed to initiate eye organ formation. It is interesting to consider these models in light of the fact that *map3k1(RNAi)* regenerating trunk fragments initially succeed at regeneration of new eyes at a normal location and only subsequently form ectopic eyes posteriorly to the first set of eyes. Given that *map3k1* transcription is induced early in regeneration, these effects could potentially be explained by differences in the effectiveness of RNAi knockdown in early versus late regeneration. Alternatively, the role of *map3k1* may be more specific to the conditions common to late blastema formation and homeostasis, in which migratory eye progenitors must travel relatively longer distances to reach their destinations compared to eyes normally formed in the first 2-3 days of blastema formation.

Whole-body regeneration likely involves a substantial component of progenitor sorting and migration. In planarians, progenitors for regionalized tissues like eyes and pharynx are specified in broad domains (6, 52). In addition, progenitors for tissues present throughout the body, such as epidermis, gut, and muscle, lack consistent spatial segregation from each other (53). Therefore, controlling differentiation of migratory progenitors to appropriate locations is likely critical for enabling appropriate regeneration. While integrins and snail-family transcription factors are important for neoblast migration and appropriate targeting of differentiating cells (54–56), and neoblast migration can be coupled to DNA damage control (57), the signals specifying target locations and also mechanisms to prevent premature differentiation are not well known. Body-wide patterning signals play an important role in determining the target locations of eye regeneration, yet *map3k1* likely controls a distinct process that prevents migratory progenitors from differentiating too early prior to arriving at their destination. We suggest that regeneration involves the integration of positional information to define progenitor domains, signaling that targets progenitors to particular locations, and systems that can sustain the undifferentiated state during migration. Our study identifies *map3k1* as a critical regulator of terminal differentiation in the eye progenitors used for organ regeneration.

## Methods and Materials

### Experimental model

Asexual *Schmidtea mediterranea* (CIW4 strain) were cultured in 1x Montjuic salts at 18-20°C. Animals were fed pureed calf liver once a week and cleaned at least once every week for maintenance. Animals were starved for at least 7 days before experiments.

### RNA interference (RNAi)

RNAi was performed by feeding dsRNA to animals every 2-3 days for the designated length of the experiment. Double stranded RNA (dsRNA) were produced by *in vitro* synthesis reactions (NxGen, Lucigen) using primers listed in Table S1. Control dsRNA was produced from *Caenorhabditis elegans unc-22*, a gene sequence not found in the planarian genome. Liver mixtures contained 20% dsRNA, 5% food coloring dye, and 75% pureed calf liver. For homeostatic RNAi treatments, animals were fixed 4 days after the last feeding. For RNAi treatments of regenerating worms, animals were amputated 1 day after the final feeding and fixed at the indicated times. Unless otherwise noted in the text, RNAi dose schedules were performed by 6 dsRNA feedings over 2 weeks. For double-RNAi experiments, single-gene RNAi comparisons to double-RNAi conditions involved mixing equal amounts of competing control dsRNA (targeting *C. elegans unc-22*) along with the experimental dsRNA to ensure the same amount of overall dsRNA was present across conditions.

### Fluorescence in situ hybridization (FISH) and immunostaining

FISH protocol used were performed as described in previous work (58). Briefly, animals were fixed in 7.5% N-Acetyl- cysteine in PBS (w/v) followed by 4% formaldehyde in PBSTx (1X PBS+0.3% Triton-X100, v/v), washed in PBSTx and stored in methanol prior to bleaching for 2-4 hours in a solution of 5% deionized formamide (v/v) and 0.36% hydrogen peroxide (v/v) in 1XSSC. Digoxigenin- or fluorescein-labeled riboprobes were synthesized as described in previous work (58) and detected with anti-digoxigenin-POD or anti-fluorescein-POD antibodies (1:2000, Roche/Sigma-Aldrich) blocked with 10% heat-inactivated Horse serum and 10% Western Blotting Blocking Reagent (Roche). Tyramide amplification was performed by depositing 0.2% rhodamine-tyramide or fluorescein-tyramide and 0.1% 4-iodophenylboronic acid (in dimethylformamide) in TSA buffer (2M NaCl, 0.1M Boric acid, pH 8.5). For double FISH, the enzymatic activity of tyramide reactions were stopped in 100mM sodium azide in TNTx. Nuclei were stained using 1:1000 Hoechst 33342 (Invitrogen) in TNTx. For immunostainings, animals were fixed in 4% formaldehyde as previously described (58), blocked with 5% heat-inactivated Horse Serum in PBSTx, incubated in goat anti-mouse HRP (1:300) to detect labeling with mouse anti-ARRESTIN (1:10,000), before tyramide amplification. Unless otherwise indicated, riboprobes were generated by PCR using primers listed in Table S1. Other riboprobes were as previously described: *GluR/gpas* (21, 30), *ChAT* (cholinergic neurons expressing choline acetyltransferase) (59), *cintillo* (60), *ovo*, *opsin*, *tyrosinase*, (6, 7), *FoxD* (61), and *ndl5* (20).

### Image Acquisition

Live animals were imaged on either a LeicaMZ125 dissecting microscope with a LeciaDFC295 camera or a LeicaS6D dissecting microscope with a flexacamC3 camera. Stained animals were imaged with a Stellaris 5 laser-scanning confocal microscope. Adjustments to brightness and contrast were made using FIJI/ImageJ or Adobe Photoshop.

### Primer Design

Primers for dsRNA and riboprobes are listed in Table S1.

### Quantification and Statistical Analysis

Stained animals were imaged using a Leica Stellaris 5 laser-scanning confocal microscope. For analysis of PCG expression domains in Figure 2, maximum projection images were analyzed in ImageJ/Fiji to measure body length, body area, and manual scoring of the extent of PCG expression. For body length, animals were measured from the head tip to the end of the tail. PCG domain measurements were conducted done by measuring the most anterior to most posterior expression for each gene. *cintillo^+^* cells in Figure 2 were counted manually and normalized to the area of the animal as determined by Hoechst staining. To quantify undifferentiated eye progenitors in Figure 6, *ovo^+^*cells located outside of eyes and also lacking *opsin/tyrosinase* expression were manually counted from maximum projections of animal head regions imaged using a 20x objective on a Leica Stellaris 5 laser-scanning confocal microscope. Quantification of *map3k1^-^ovo^+^* and *map3k1^+^ovo^+^* cells in Figure S4D was conducted through manual scoring of confocal z-stacks acquired using a 40X glycerol-immersion objective. Sample sizes for each experiment are indicated in the legends and/or through plotting individual datapoints for each experiment. Statistical tests are indicated in each figure legend and were conducted in Microsoft Excel, boxplots were generated using BoxPlotR, stacked bargraphs were generated in Microsoft Excel or Datawrapper, and 2D dot plots were generated in R.

### Counting *opsin^+^* cells within planarian eyes

Confocal z-stacks were obtained on a Leica Stellaris 5 laser-scanning confocal microscope using a 40x glycerol objective and 0.3-micron slice size. For each sample, individual left and right sides of the head were imaged using equivalent laser and gain settings, and z-stack size chosen to capture the entire depth of all eye cells. *opsin*^+^ cell numbers were quantified using Stardist to segment nuclei (probThresh=0.60, nmsThresh=0.5) and counting nuclei ROIs whose median opsin-channel fluorescence exceeded an empirically defined threshold of detection obtained from Otsu thresholding slices that contained target cells, measured in ImageJ. *opsin*^+^ cell counts from individual slices taken every 5-microns were summed across a z-stack for each sample. Total *opsin*^+^ cells were compared across control (n=8) and *map3k1*(RNAi) (n=8) treatments using an unpaired 2-tailed t-test. The analysis was performed across a range of threshold settings (+/- 10% pixel intensity of threshold) and slice widths (from 2.5 to 10 microns), and similar differences to *opsin*^+^ cell numbers were measured in each case across the treatment types. A Jaccard similarity index/coefficient (JSI, Intersection over Union) was calculated by randomly selecting 6 annotated z-stacks collected from control and *map3k1(RNAi)* worms which were used in the calculating the estimated number of *opsin^+^* cells, and manually categorizing cells as true positives, false negatives, and false positives (Table S2). The average JSI was calculated for control and *map3k1* RNAi samples separately, and an unpaired 2-tailed t-test was performed to determine whether automated cell segmentation and counting efficiency varied between treatment types.

### Mapping relative positions of eyes in fixed and live animals

In Figures 5 and 8, images of live or stained animals were overlaid on a grid, which normalized to the inter-eye distance such that the left and right pigment cup cells were positioned at (-1,0) and (1,0) respectively. For analysis of ectopic eye phenotypes, the pigment cup of each ectopic eye was marked, and the distance from the original eyes was measured using WebPlotDigitizer (https://apps.automeris.io/wpd/), and then normalized in units equal to one half the inter-eye distance. A similar procedure was used to map the relative positions of *ovo+* progenitor cells in images of fixed and stained animals. The relative coordinates of each eye or *ovo+* eye progenitor were extracted and plotted using ggplot2 in R.

## Acknowledgements

We thank members of the Petersen lab for critical comments and thoughtful discussion. We thank R. Zayas for the kind gift of the planarian anti-ARRESTIN antibody.

## Author Contributions

Katherine C. Lo: Conceptualization, data curation, formal analysis, investigation, methodology, validation, visualization, writing- original draft & preparation.

Christian P. Petersen: Conceptualization, funding acquisition, methodology, resources, supervision, writing- original draft & preparation

## Supplementary Figure Legends

**Figure S1.**
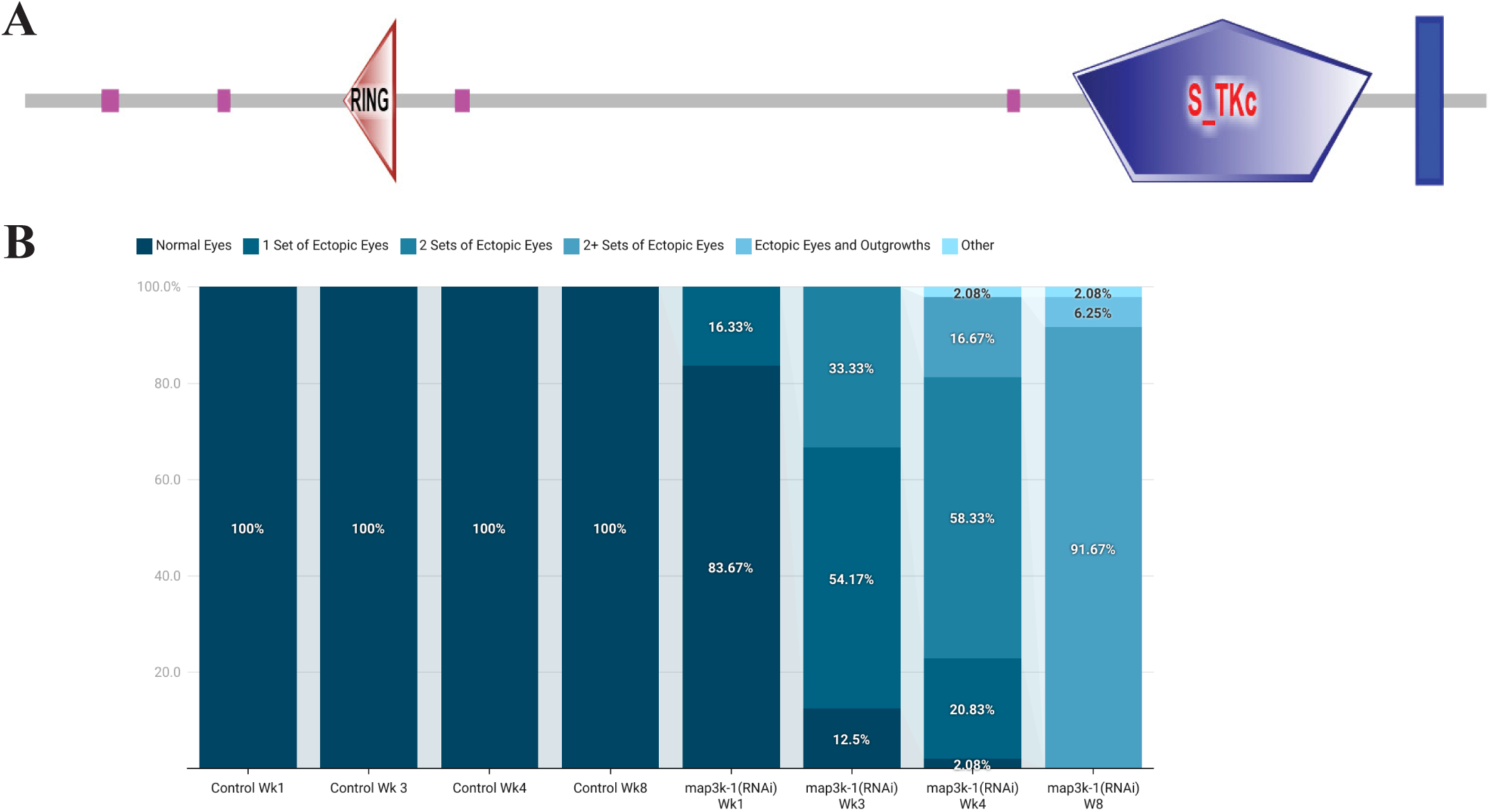
(A) Domain structure of dd_Smed_v6_5198_0_1 (*map3k1)* containing a RING (E-value= 0.0255) domain, a serine/threonine kinase domain (E-value=1.16e-69) characteristic of MAPKs, and a transmembrane region. (B) Stacked bar graph quantifying the number of ectopic eyes in control (n=46) versus *map3k1(RNAi)* (n=48) animals over 8 weeks of RNAi showing that *map3k1* inhibition caused ectopic eyes to continue forming over time.

**Figure S2.**
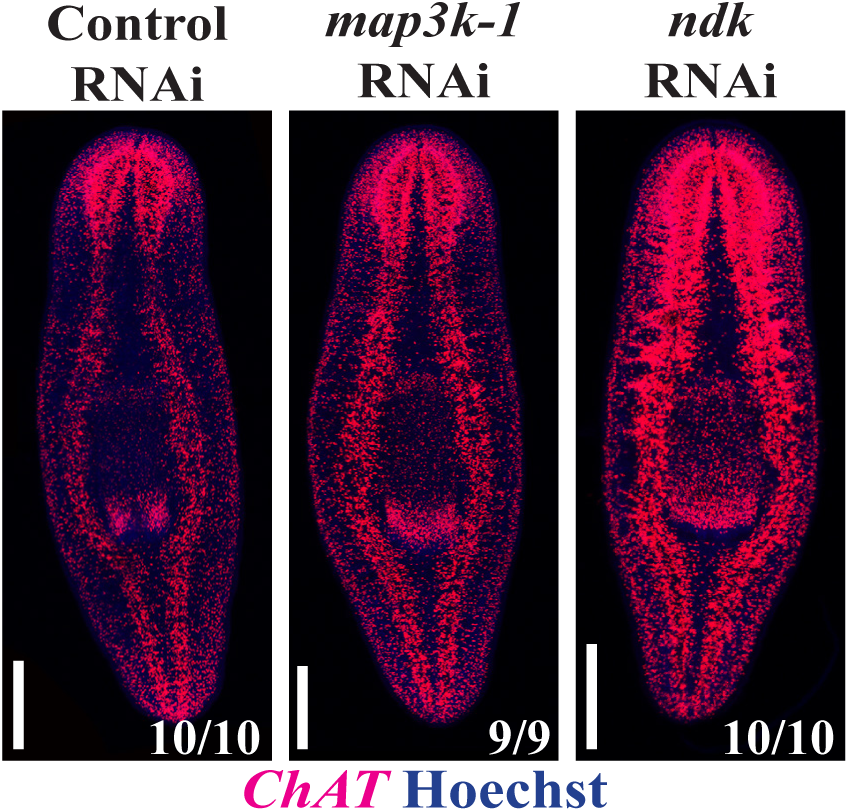
Homeostatic animals fixed after 4 weeks of RNAi were stained with *ChAT* to detect cholinergic neurons. *map3k1* RNAi did not increase *ChAT^+^* neuron staining, compared to the ectopic *ChAT^+^* brain branches that formed after *ndk* RNAi. n≥9. Scale bars, 300μm.

**Figure S3.**
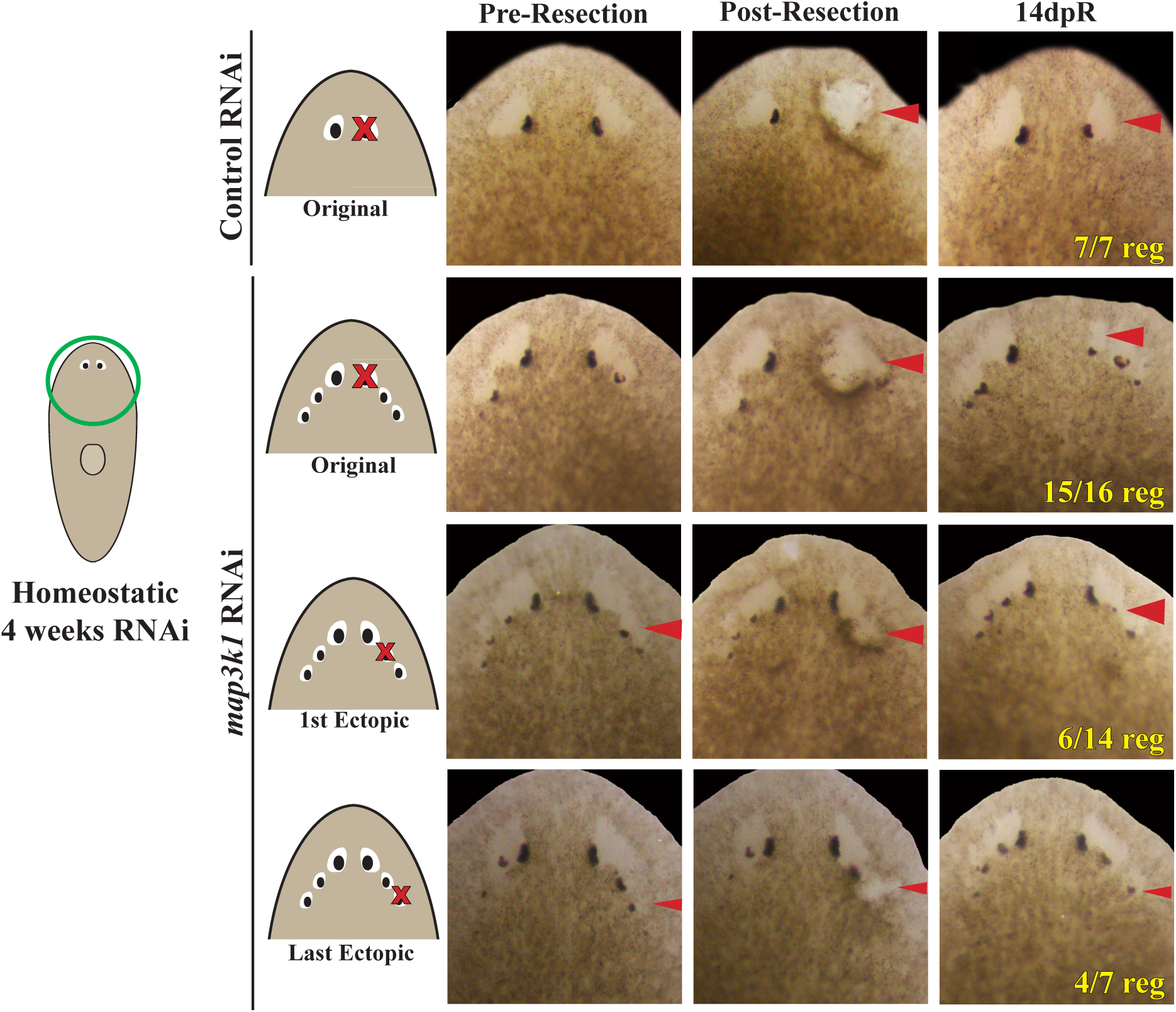
To test whether *map3k1* could control eye placement and/or a subset of head patterning, animals were fed with either control or *map3k1* dsRNA for 4 weeks before undergoing eye resections at different positions indicated by the cartoons. Individual live animals were imaged before, immediately after (post-resection), and 14 days post-surgical eye resection (14dpR) to track whether eye regeneration subsequently occurred. *map3k1(RNAi)* animals regenerated their original eyes at a high frequency (15/16). Ectopic eyes from these animals were also capable of regeneration, though at lower frequencies. Removal of either the anterior-most ectopic eyes (6/14 eyes regenerated, “1^st^ ectopic”) or the posterior-most ectopic eyes (4/7 eyes regenerated, “last ectopic”) could result in regeneration from the original eye. Sample size, n≥7 animals in each condition.

**Figure S4.**
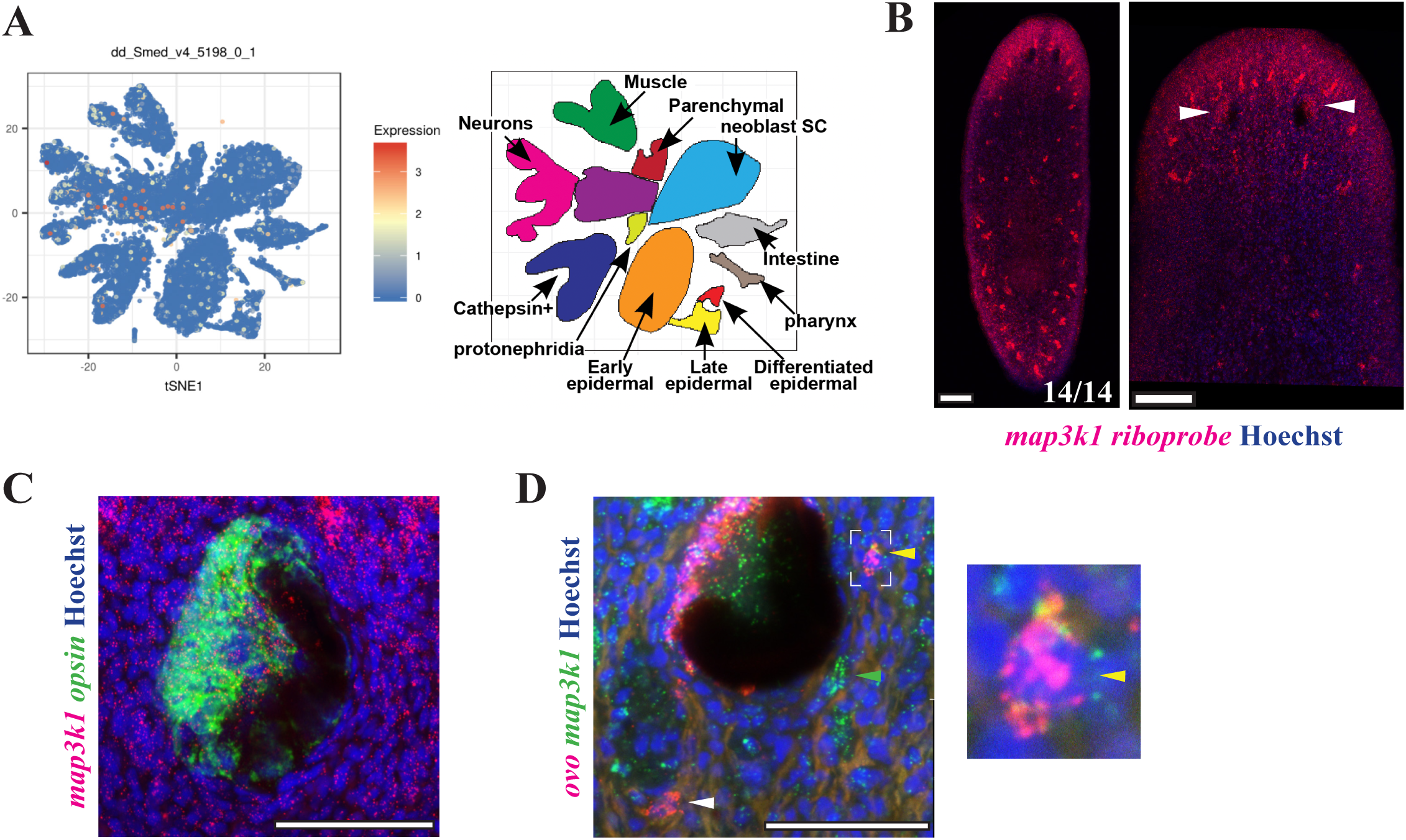
(A) Single-cell RNA sequencing expression profile of *map3k1* during injury as published from a prior study (31). *map3k1* is expressed broadly and in many different tissues, but has enriched expressed in the muscle, neural, and gut clusters. (B) Maximum projection images of *map3k1* expression in homeostatic worms as detected by FISH show *map3k1* is expressed broadly throughout the body. Right panel, image showing *map3k1* expression in the head and low levels of *map3k1* expression in the eyes (arrows). Sample size, n=14 animals. Scale bar, 100μm. (C) Maximum projection images of *map3k1* and *opsin* expression in the eye show some expression of *map3k1* in *opsin* expressing cells (4/4 animals). Scale bar, 50μm. (D) Double-FISH detecting *ovo* and *map3k1* expression. Some ovo*^+^* cells expressed low levels of *map3k1* (yellow arrow, 22/36 cells counted over 5 intact animals) while others did not have *map3k1* expression (white arrow, 14/36 cells over 5 animals). *map3k1* is also broadly expressed, so other unknown *map3k1^+^* but *ovo^-^* cells were identifiable (green arrow). Scale bar, 50μm.

**Table S1.**
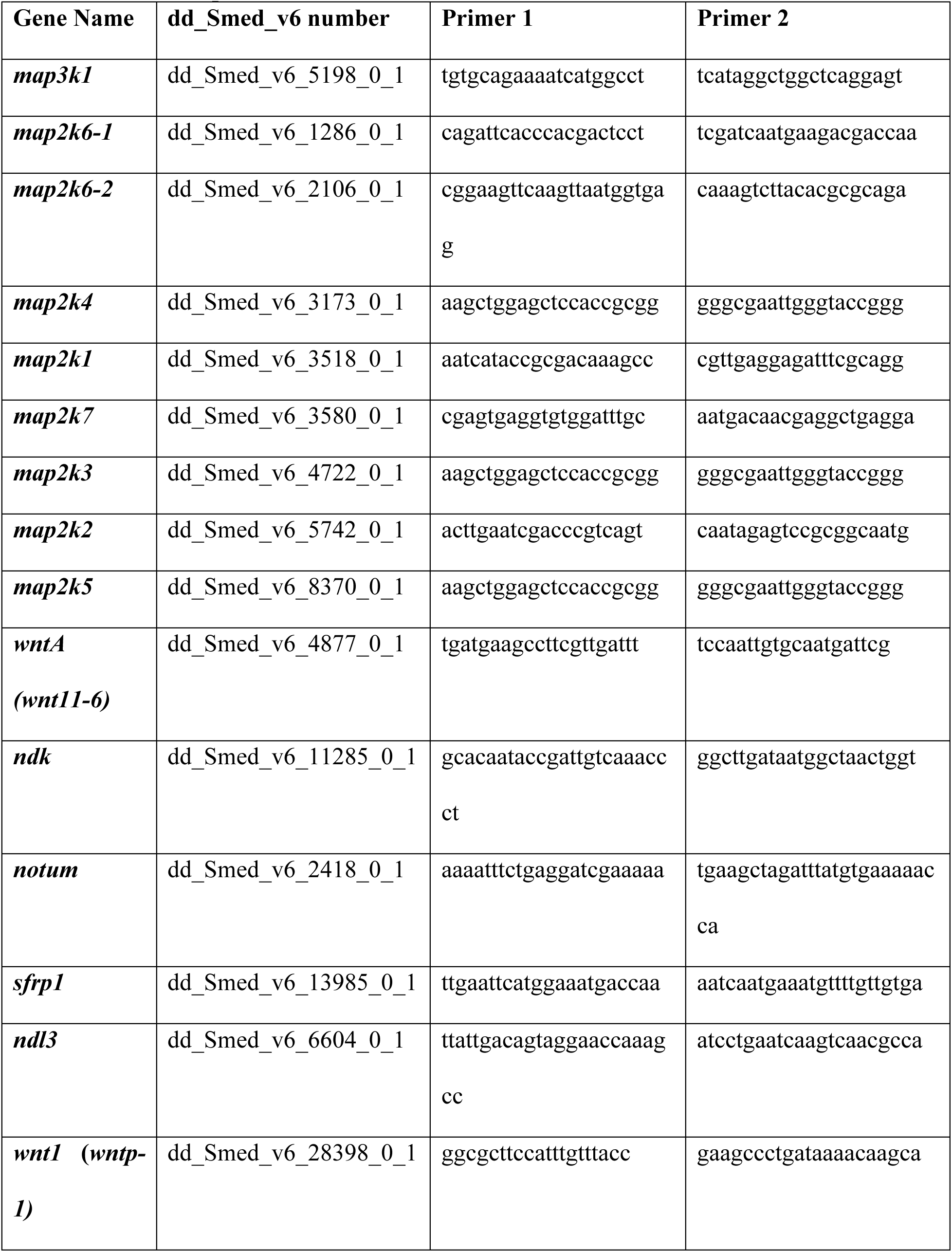
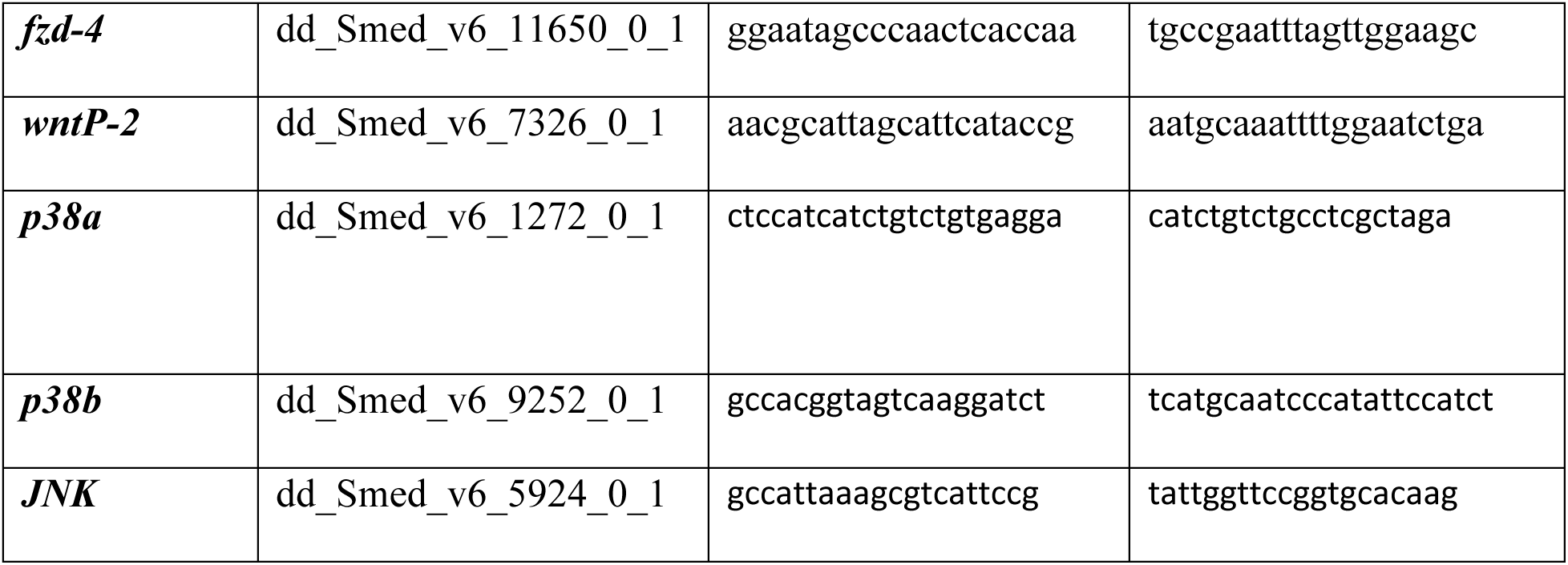
Primer sequences.

**Table S2.** Manual Cell Count Values for Nuclei Segmentation.

| **Image Name** | **True**<br>**Positives**<br>**(TP)** | **False**<br>**Negative**<br>**(FN)** | **False**<br>**Positive**<br>**(FP)** | **Total Cells**<br>**(TP+FN+FP)** | **Jaccard**<br>**Index** |
| --- | --- | --- | --- | --- | --- |
| *Control(RNAi)* Sample 1 | 9 | 5 | 0 | 14 | 0.642857 |
| *Control(RNAi)* Sample 2 | 5 | 2 | 1 | 8 | 0.625 |
| *Control(RNAi)* Sample 3 | 11 | 10 | 2 | 23 | 0.478261 |
| *Control(RNAi)* Sample 4 | 10 | 11 | 1 | 22 | 0.454545 |
| *Control(RNAi)* Sample 5 | 8 | 6 | 4 | 18 | 0.444444 |
| *Control(RNAi)* Sample 6 | 10 | 2 | 3 | 15 | 0.666667 |
| *map3k1(RNAi)* Sample 1 | 34 | 10 | 11 | 55 | 0.618182 |
| *map3k1(RNAi)* Sample 2 | 19 | 15 | 0 | 34 | 0.558824 |
| *map3k1(RNAi)* Sample 3 | 19 | 20 | 4 | 43 | 0.44186 |
| *map3k1(RNAi)* Sample 4 | 33 | 11 | 14 | 58 | 0.568966 |
| *map3k1(RNAi)* Sample 5 | 9 | 21 | 0 | 30 | 0.3 |
| *map3k1(RNAi)* Sample 6 | 19 | 21 | 3 | 43 | 0.44186 |

